# Performance of a phylogenetic independent contrast method and an improved pairwise comparison under different scenarios of trait evolution after speciation and duplication

**DOI:** 10.1101/2020.09.25.313353

**Authors:** Tina Begum, Martha Liliana Serrano-Serrano, Marc Robinson-Rechavi

## Abstract

1. Despite the importance of gene function to evolutionary biology, the applicability of comparative methods to gene function is poorly known. A specific case which has crystalized methodological questions is the “ortholog conjecture”, the hypothesis that function evolves faster after duplication (i.e., in paralogs), and conversely is conserved between orthologs. Since the mode of functional evolution after duplication is not well known, we investigate under which reasonable evolutionary scenarios phylogenetic independent contrasts or pairwise comparisons can recover a putative signal of different functional evolution between orthologs and paralogs.
2. We investigate three different simulation models, which represent reasonable but simplified hypotheses about the evolution of a gene function trait. These are time dependent trait acceleration, correlated changes in rates of both sequence and trait evolution, and asymmetric trait jump. For each model we tested phylogenetic independent contrasts and an improved pairwise comparison method which accounts for interactions between events and node age.
3. Both approaches lose power to detect the trend of functional evolution when the functional trait accelerates for a long time following duplication for trees with many duplications, with better power of phylogenetic contrasts under intermediate scenarios. Concomitant increase in evolutionary rates of sequence and of trait after duplication can lead to both an incorrect rejection of the null under null simulations of trait evolution, and a false rejection of the ortholog conjecture under ortholog conjecture simulations, by phylogenetic independent contrasts. Improved pairwise comparisons are robust to this bias. Both approaches perform equally well under rapid shifts in traits.
4. Considering our ignorance of gene function evolution, and the potential for bias under simple models, we recommend methodological pluralism in studying gene family evolution. Functional phylogenomics is complex and results supported by only one method should be treated with caution.

## Introduction

In comparative biology, pairwise comparisons of terminal taxa of genes or of species are commonly used to detect phenotypic or morphological or character associations (Felsenstein 1985, Maddison 2000; Martins & Garland 1991; Read & Nee 1995). This pairwise approach can handle sparse data and poorly resolved phylogenies, and relies on relatively few assumptions, compared to other methods that use reconstructed ancestral states, stochastic models of evolution, or phylogenetic branch lengths (Felsenstein 1985; Read & Nee 1995; Purvis & Bromham 1997; Garland *et al*. 1992; Maddison 2000). A potential issue is that multiple pairwise comparisons of data across the same branch of a tree pseudo-replicate data, and overlook phylogenetic relatedness of species (Pagel 1994; Blomberg et al. 2003; Dunn *et al*. 2018). The phylogenetic independent contrasts (PIC) method solves these issues by taking into account the evolutionary history of species (Felsenstein 1985; Grafen 1989; Cooper *et al*. 2016; Dunn *et al*. 2018). This approach needs well-resolved phylogenies. Moreover, it is sensitive to model assumptions (e.g. known branch lengths of the phylogeny, or Brownian character evolution), and may lead to erroneous interpretations if these assumptions are violated (Garland *et al*. 1992; Maddison 2000; DÌaz-Uriarte & Garland 1998; Freckleton & Harvey 2006; Cooper *et al*. 2016). Most comparative functional genomic studies still rely on pairwise comparisons, although some do use PIC.

Applications of both approaches to test the “ortholog conjecture” model, i.e. that there are larger functional differences between paralogs (homologous genes diverging since a duplication) than between orthologs (homologous genes diverging since a speciation), have shown how methodological differences can lead to different results concerning the evolution of gene function (e.g., Nehrt *et al*. 2011; Kryuchkova-Mostacci & Robinson-Rechavi 2016; Dunn *et al*. 2018; Begum & Robinson-Rechavi 2020). Despite these conflicting empirical results, the theoretical expectations of these methods under different models of evolution of gene function have not been studied.

Two common features of most observations of gene evolution are asymmetry of rate changes between phylogenetic branches, and that sequence rates are variable (i.e., there is no molecular clock). Both of these can be expected to affect testing of the ortholog conjecture. Functional traits evolve asymmetrically after duplication (Gu *et al*. 2005; Kim & Yi 2006; Cusack & Wolfe 2007; Studer & Robinson-Rechavi 2009). At the sequence level (e.g., dN/dS), there appears to be both an acceleration following gene duplication, and asymmetry of this acceleration (Conant & Wagner 2003; Brunet *et al*. 2006; Kim & Yi 2006; Scannell & Wolfe 2008; Studer & Robinson-Rechavi 2009; Panchin *et al*. 2010; Pegueroles *et al*. 2013; Pich & Kondrashov 2014; Holland *et al*. 2017). In yeast, Gu *et al*. (2005) reported asymmetric trait acceleration in a time-dependent fashion shortly after duplication, with less dramatically acceleration in sequence evolutionary rates. Trait evolutionary rates are also variable in phylogenies. For example, there can be a period of rapid trait shifts or jumps according to changes in the fitness landscape (e.g. ecological opportunities) following a speciation event (Bokma 2008; Duchen *et al*. 2017; Landis *et al*. 2013; Simpson 1944). Such a jump in traits, rather than a change in the evolutionary rates, might also affect trait evolution after gene duplication. The complexity of evolution after duplication led us to enquire under which reasonable simple evolutionary scenarios after gene duplication we can recover a signal of different functional evolution between types of events in gene trees.

To address this question, we consider the ortholog conjecture as a test case and we use a gene trait with a bound limit of 0 to 1; we call it *τ*, like the tissue-specificity score used in previous studies (Yanai *et al*. 2005; Kryuchkova-Mostacci & Robinson-Rechavi 2016; Dunn *et al*. 2018). An advantage of simulations is that we have perfect knowledge of the original time trees (Fig 1A), which we never know for empirical data. With empirical data, we obtain “Substitution trees” (Fig 1B), which are time calibrated using the speciation time points to generate pseudo time trees, i.e. “Calibrated time trees” (Fig 1C), for hypothesis testing. We compared the performances of PIC and of an improved pairwise method using three simple simulation models with different parameters of divergence following speciations and duplications. We used two sets of simulated trees: i) trees with arbitrary parameter values and different proportions of duplications, which we call “pure simulated” trees, and ii) trees simulated using parameter values (speciation rates, extinction rates, number of tips, and proportions of internal node events) from calibrated dichotomous empirical gene trees. The first allow to explore parameter space in an unbiased way, while the latter allow to explore more realistic scenarios (S1 Table) but can be affected by bias of duplication branch lengths (Begum & Robinson-Rechavi 2020).

**Fig 1:**
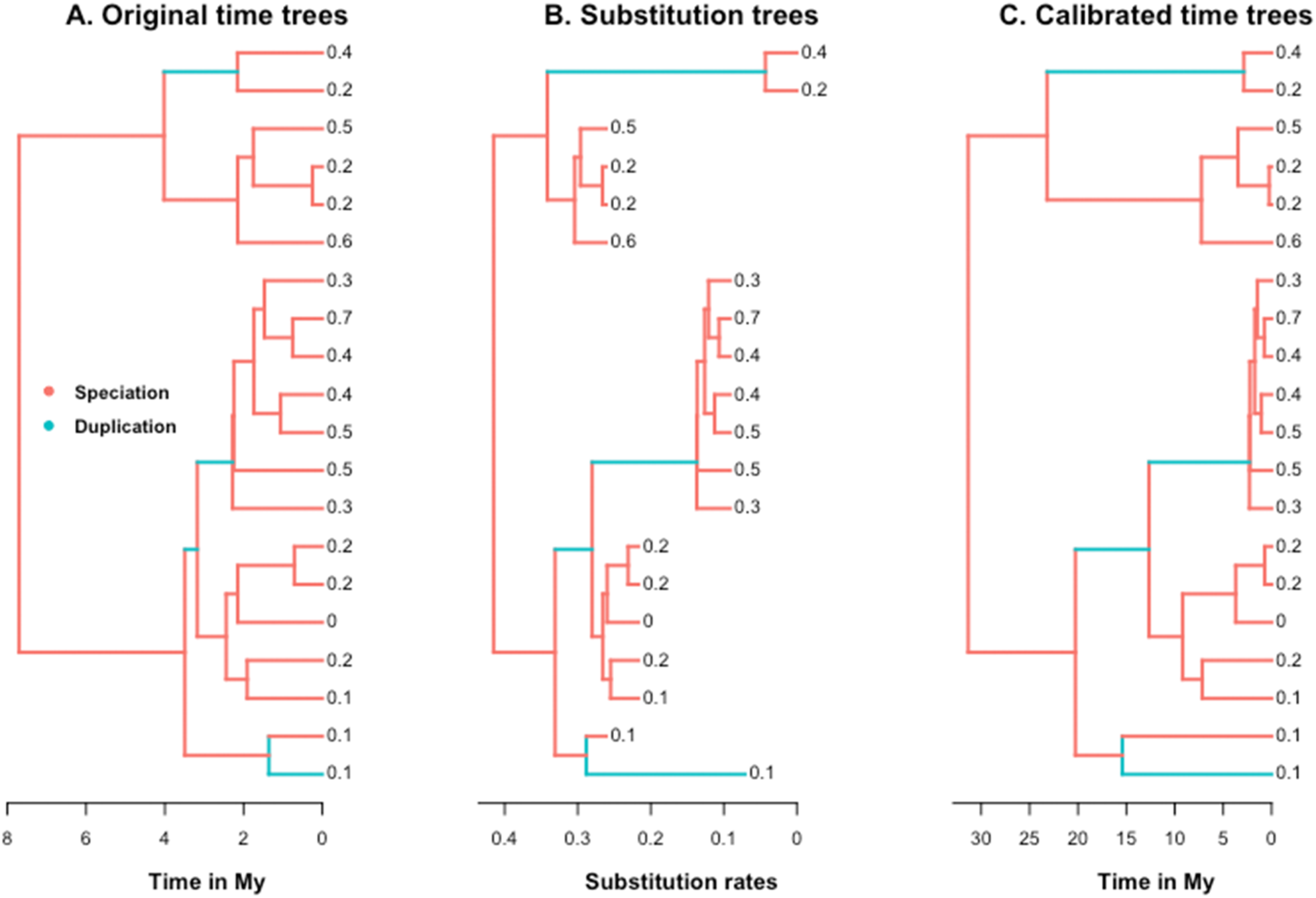
An example of simulated tree models used in this study. (A) Original (true) time tree, (B) substitution rates tree, and (C) pseudo time tree inferred from (B), with simulated trait values at tips. We used a simulated tree with 20 tips for illustration. Branches following gene duplication are asymmetrically painted using the phytools R package (Revell 2012) to introduce different sequence evolutionary rates to those branches. This demonstrates how the scale (i.e. the branch lengths) of the calibrated tree can differ from the original time tree.

## Materials and Methods

### Simulated gene trees

We simulated ultrametric trees (n = 10000), each with 100 tips generated under a pure birth-death model using the TreeSim R package (Stadler 2011) with a speciation rate of 0.4, and an extinction rate of 0.1 for pure simulated trees (S1 Table). Since our study aims to mimic simple evolutionary scenarios (test model details in Supplementary text), where standard trait evolutionary models are frequently applied on comparatively smaller phylogenies (Chira & Thomas 2016), we limited our tree size to100 tips. We call these simulated time trees “Original time trees” (Fig 1A), with the branch lengths corresponding to units of time (i.e. Million Years – My). For consistency, we used the same set of 10000 original time trees for all further analyses.

### Annotation of internal node events

We annotated internal node events as “speciation” or “duplication” so that each pure simulated tree had at least one speciation and one duplication node events. We considered 3 proportions of duplication events (number of duplication events / total number of internal nodes): 0.2, 0.5, and 0.8 (S1 Fig, S1 Table). We first randomly annotated “duplication” events to internal nodes based on the proportion of duplication events we considered for the tree set. We then assigned “speciation” events to the rest of the nodes of the tree.

From empirical vertebrate data, we find the median proportion of duplications events in a range of 0.1 - 0.2 (Dunn *et al*. 2018; Begum & Robinson-Rechavi 2020). Therefore, we used the pure simulated tree set with a proportion 0.2 of duplication events (S1 Table) in all our main text analyses to compare the performances of different approaches in testing the ortholog conjecture on different simulation models. To verify that our results did not depend on the number of duplication events in a tree, we reproduced results using proportions of duplications of 0.5 and 0.8 (supporting materials).

### Simulation of trait

In this study, we considered a trait with a range between 0 to 1. We call it, *τ* like the tissue-specificity score used in several previous studies of the ortholog conjecture (Yanai *et al*. 2005; Kryuchkova-Mostacci & Robinson-Rechavi 2016; Dunn *et al*. 2018). For the sake of simplicity, we only used a Brownian model (BM) of trait evolution:

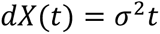

where *dX*(*t*) quantifies the change in trait, and the Brownian variance (*σ*^2^ paremeter describing the rate at which taxa diverge from each other through time *t* (Cavalli-sforza & Edwards, 1967; Harmon *et al*. 2010; Chira & Thomas 2016). We used FastBM function of the Phytools R package (Revell 2012) to simulate the trait *τ* on each original time tree. The same trait values were used for the corresponding substitution rate trees (described later), and for the pseudo time calibrated trees (described later) for further analyses with phylogenetic method.

### Generation of calibrated pseudo time trees

We used the NELSI R package (Ho *et al*. 2015), and modified the original *simulate.rates* function to *simulate.rates_heterogeneous* to introduce different sequence evolutionary rates (nucleotide substitutions/site/My) to the branches of different node events with the aid of ‘discrete’ multi-rates clock model. To avoid bias in our inference, we used the same sequence evolutionary rates, K, as used for simulation of the trait *τ* in a null scenario. These trees are “Substitution trees” (Fig 1B), where the branch lengths of each tree representing substitution rates. In these phylograms, the branch lengths are proportional to the number of mutations accumulated along the branches, and thus changes in branch lengths should have an influence on both the nonsynonymous and synonymous substitution rates.

We used the *chronos* function of the APE R package (Paradis *et al*. 2004) to generate calibrated time trees (Fig 1C). For such time calibration, we normally have access to only a few speciation time points (focal speciation nodes) for which we have external references (e.g. fossil data) (Dunn *et al*. 2018; Begum & Robinson-Rechavi 2020). However, due to simulated tree structure, very few trees passed time calibration step when we fix focal speciation nodes, and their ages across all trees. Hence, we used the ages of all the speciation nodes of a tree to time calibrate them. The scale of the original tree drastically changes after calibration if we do not fix the age of the root node. It may impact the inference of phylogenetic method due to its branch length dependence. Hence, we also used the age of the root of each simulated original time tree to maintain the scale after time calibration.

### Simulations with empirical data parameters

We downloaded 60447 gene trees from ENSEMBL Compara v.100 (Herrero *et al*. 2016), and annotated tips with precomputed tissue specificity data, *τ* covering six organs of 21 vertebrate species from Fukushima and Pollock (2020). Tips with missing trait data were pruned. When two speciation nodes had the same clade names at different node depths of a tree, the older speciation event was edited to “NA” to prevent failure of time calibration (Dunn *et al*. 2018). This led us to obtain 13647 calibrated empirical gene trees.

To simulate original time trees based on empirical data parameters, we used only dichotomous trees, i.e., 6953 out of 13647. This helps us to avoid over-estimated speciation and extinction rates using the *birthdeath* function of the APE R package (Paradis *et al*. 2004). The estimated rate parameters for each tree, including tip number, were used to simulate 6953 original time trees using TreeSim (Stadler 2011). Since the actual tree topology changes after simulation, we randomly assigned node events for each tree, while maintaining the empirical event proportions (S1 Table).

In this case, we estimated trait evolutionary rates for each calibrated time tree based on empirical *τ* of tips following a BM using the *fitContinuous* function (Pennell *et al*. 2014). We used them to simulate trait for our 3 test models.

### Phylogenetic analyses

For each internal node (inode) of a gene tree, we calculated the PIC of simulated trait *τ* from the APE R package using the following formula:

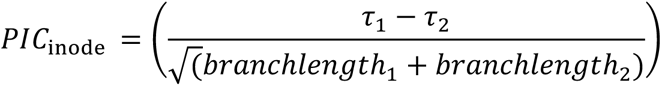

where PIC_inode_ is the node of which PIC is to be computed, *τ*_1_, and *τ*_2_ are the trait values of daughters of the corresponding node. The branch lengths of the daughters of corresponding node are represented by branchlength_1_, and branchlength_2_, respectively (Felsenstein1985; Paradis *et al*. 2004; Dunn *et al*. 2018). We then contrasted the PIC of each event (speciation or duplication) using a Wilcoxon two-tailed rank test.

### Pairwise analyses

We considered all possible pairwise comparisons for each tree to obtain Pearson correlation coefficients r between *τ* values of events, as was done by Dunn et al. (2018). Few correlation points of speciation and duplication events often make it difficult to infer the statistical significance level precisely. Since most of the pure simulated trees have maximum node ages less than 20 My, we computed the correlation coefficient, r of speciation and of duplication events over each 0.05 My time intervals. To analyze simulated trees with empirical parameters, the time interval is set to 10 My since we have maximum node ages greater than 2000 My. This gives the estimates of correlation for both the events at equal time points. Previous pairwise comparisons of Kryuchkova-Mostacci and Robinson-Rechavi (2016) used a linear model (i.e. lm(R ∼ Event)) to distinguish the effects of events on correlation coefficients. In contrast, we used polynomial (linear, quadratic, or cubic) regression in our improved pairwise analyses, and used repeated 10 fold cross validation approach to choose the best fit model. The model with highest adjusted R^2^ value and with the lowest root mean square error among the linear, quadratic or cubic models was chosen as the best fit model for each case to avoid over fitting. We then compared with ANOVA the best-fit polynomial with and without the interaction terms between events and age, to obtain our improved pairwise *P* values.

### Statistical significance

For any method, we considered a result significant if *P* < *α*′. We applied a Bonferroni correction over the 148 tests that we performed overall in this study, with *α* = 0.05, thus *α*′ = 0.05 / 148 = 3.38 10^−4^.

### Code availability

For reproducibility, all scripts are available at https://github.com/tbegum/Method-matters-in-gene-family-evolution.

## Results

### Model-1: Time dependent trait acceleration model

Immediately after duplication, both duplicates might experience accelerated evolution for a period of time due to relaxed selection (Rogozin 2014). This can be represented by a simple model of acceleration of trait evolutionary rate for a given time after duplication (Fig 2). After that period of acceleration, the evolutionary rates return to the pre-duplication level. In this model, speciation does not have any impact on evolutionary rates: a speciation during the time period of the duplication-caused acceleration does not revert the gene to a “speciation” rate. To study our capacity to recover evolutionary signal under this model, we simulated with different durations of accelerated evolution, and different proportions of duplications to speciations in the gene trees (Fig 2). In this comparative study, we used an improved pairwise comparison due to the low power of the simple pairwise comparison approach used in Kryuchkova-Mostacci and Robinson-Rechavi (2016) (S2 Table).

**Fig 2:**
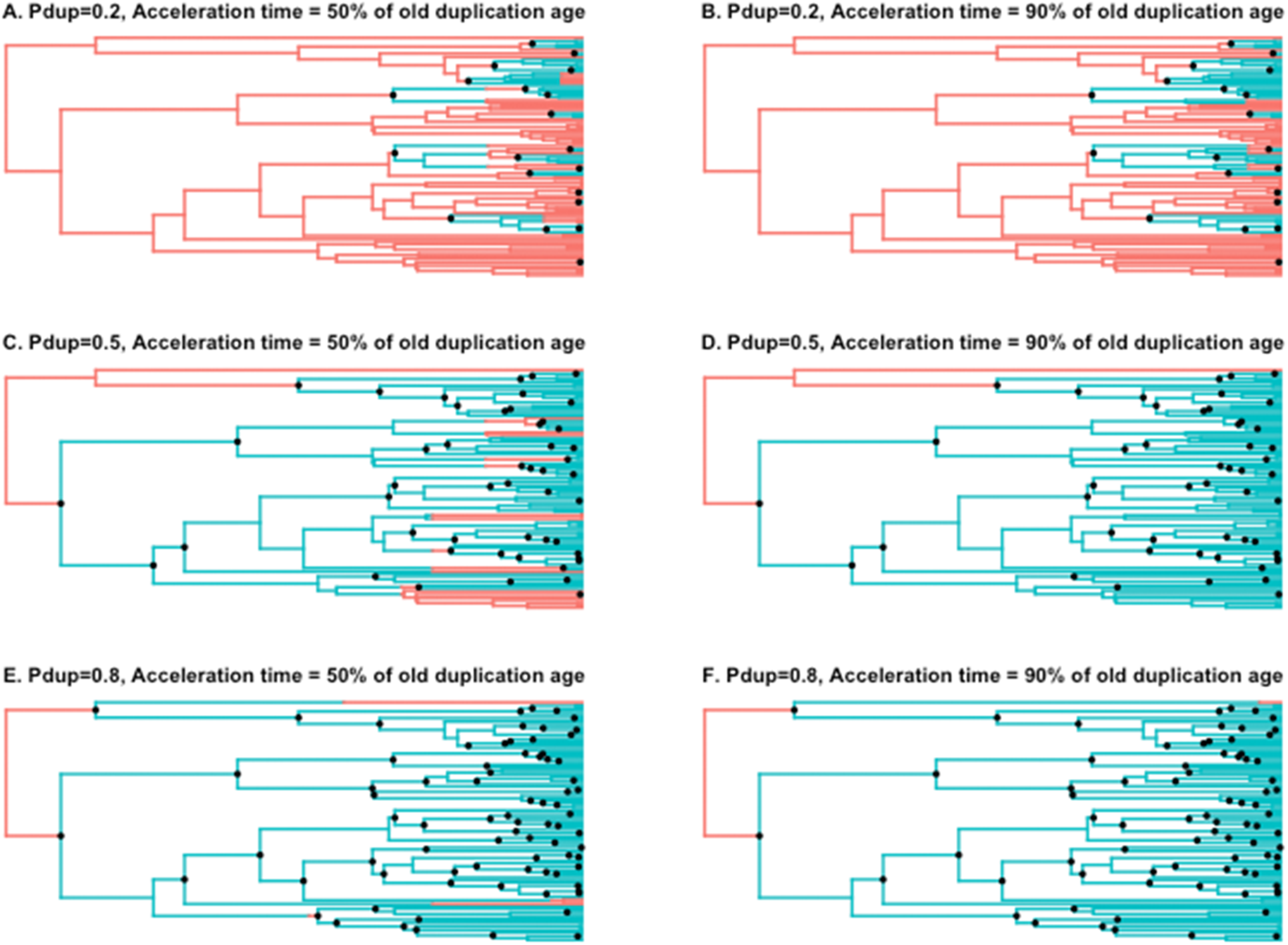
Example of time dependent trait acceleration model. We painted trees to demonstrate the effect of trait acceleration for 50% or 90% of the time since the oldest duplication, respectively. We show here a symmetrical model of trait acceleration, i.e. after duplication both the duplicates experience trait accelerations. Pdup: Proportion of duplication. Duplication nodes are indicated by black dots, while others are speciations.

First, both methods recover the proper evolutionary signal under the null, with no difference of PIC between duplication and speciation branches (Fig 3A), and no difference between ortholog and paralog pairs (Fig 3B). Second, when there are few duplications in proportion to speciations, both approaches recover the evolutionary signal, namely faster evolution of paralogs (pairwise) or of duplication branches (PIC) (Fig 3C-F, S2 Table). This is despite the acceleration lasting 50% to 90% of the time since the oldest duplication. This is possible because the rarity of duplications provides many speciation branches without acceleration (Fig 2). While the performance of both methods at recovering the evolutionary signal decreases when the proportion of duplication nodes increases, the PIC is more robust than the pairwise approach (S2 Table). With 50% of duplication nodes, only the PIC still recovers the evolutionary trend when the trait acceleration period increases. At 80% of duplications and an acceleration lasting 70% to 90% of old duplication age, neither approach can recover the signal of the ortholog conjecture (S2 Table). Under such scenarios, most speciation branches or ortholog pairs have evolved mostly or entirely under the influence of a duplication-caused acceleration, and there is no signal to recover.

**Fig 3:**
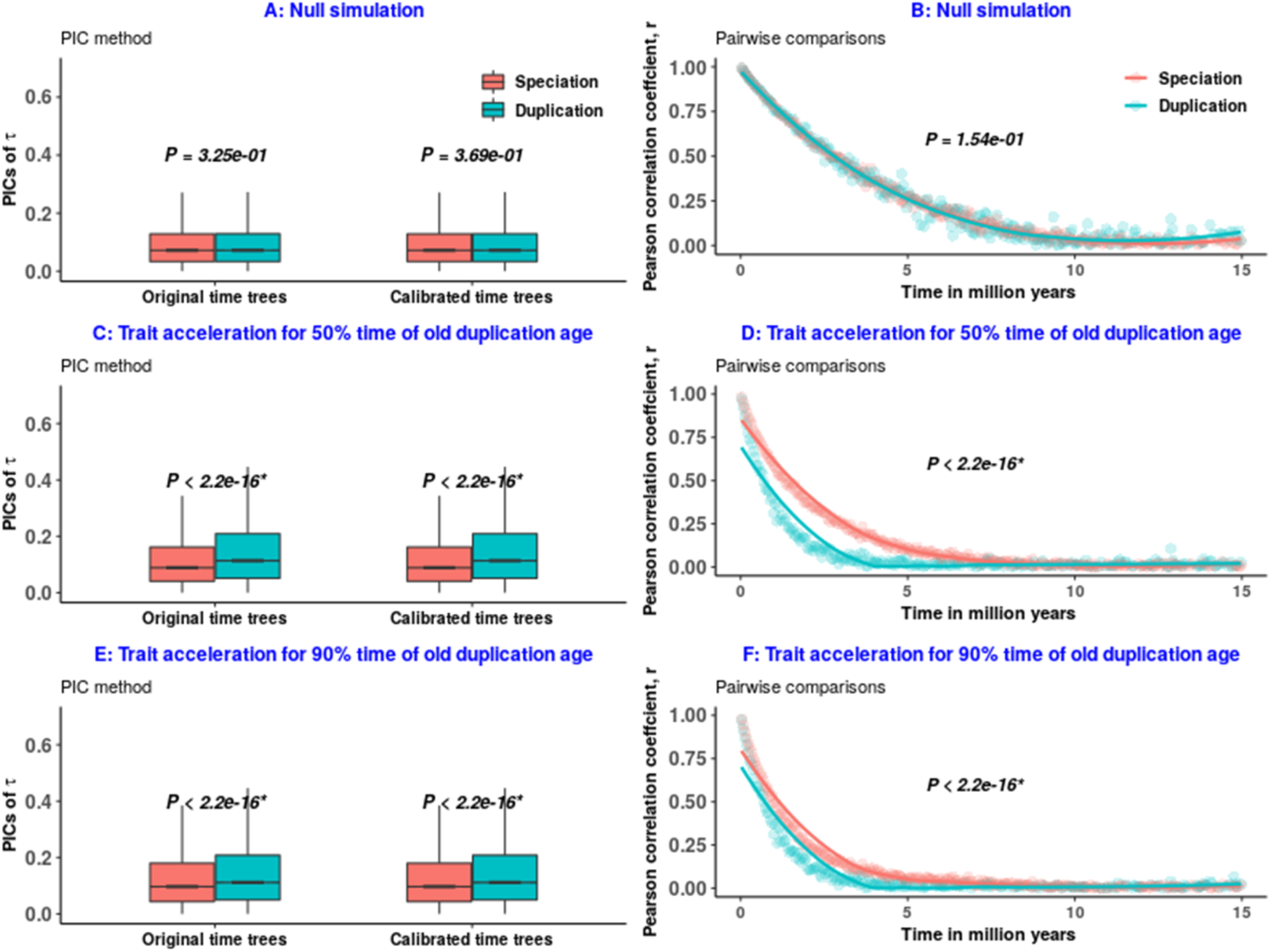
Performances of approaches in different time periods of trait accelerations since oldest duplication. Trait evolutionary rates are 5 times faster for duplication than speciation. PICs: Phylogenetic Independent Contrasts. In this symmetrical pure simulation model of 0.2 proportion of duplications, we used a cubic fit polynomial in our improved pairwise approach. *: Bonferroni-corrected significant *P* values.

We obtained similar trends for the trees simulated using empirical data parameters when we used sufficiently high trait acceleration rates following duplication (S3 Table), although the PIC then failed under the null. With only a slight acceleration (1.2 or 2 times), the phylogenetic method incorrectly rejects the null under a null simulation, and shows an opposite trend under the ortholog conjecture simulations of trait evolution. In these cases, the improved pairwise comparisons can still recover simulated trends, in many scenarios including the null, although with low power (S3 Table). The failure of PIC is probably due to the bias in branch lengths when using parameter values from calibrated empirical gene trees.

### Model-2: Rates of sequence and trait evolution model

While there is debate over the evolutionary rate of gene function after duplication, it has been shown repeatedly that there is an acceleration of sequence evolutionary rates (Conant & Wagner 2003; Gu *et al*. 2005; Brunet *et al*. 2006; Kim & Yi 2006; Cusack & Wolfe 2007; Scannell & Wolfe 2008; Studer & Robinson-Rechavi 2009; Panchin *et al*. 2010; Pegueroles *et al*. 2013; Pich & Kondrashov 2014; Holland *et al*. 2017). We explore the impact of this acceleration on the PIC and improved pairwise comparisons under different scenarios (Table 1). Because many empirical studies have found asymmetry of this acceleration of duplicate gene sequence evolution, we used an asymmetric acceleration in our simulations.

**Table 1:**
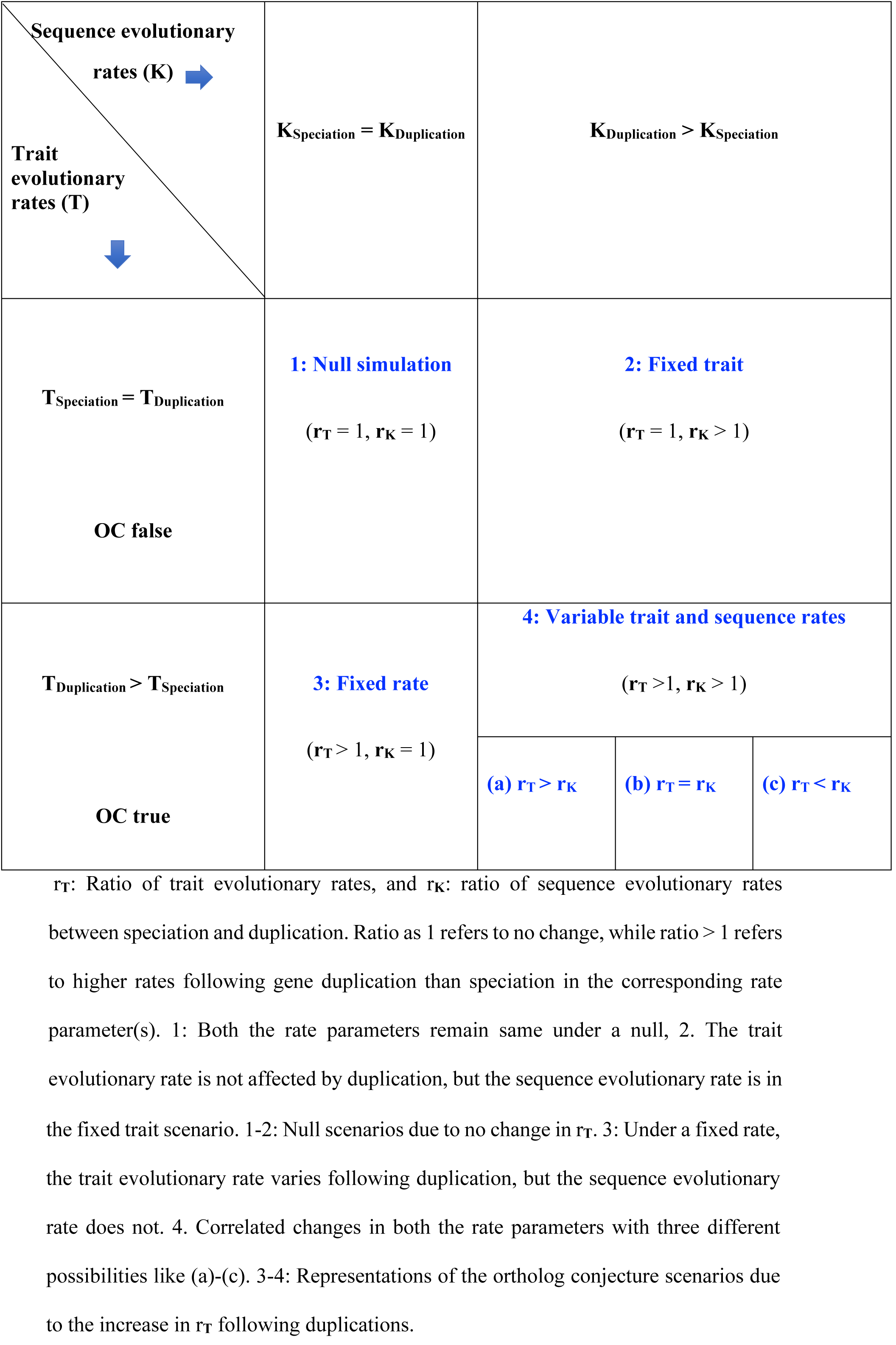
Hypothetical scenarios of the rates of sequence and trait evolution model.

**Table 2:** Summary table for 3 simulation models based on pure simulations.

| **Simulation model** | **Relevant parameters** | **Performance of PIC method** | **Performance of improved pairwise comparisons** |
| --- | --- | --- | --- |
| Time dependent trait acceleration | Trait acceleration time, and heterogeneous rates of trait evolution between events | Performs well for trees with few, and intermediate duplication events. It lacks power to detect the actual trend when trait accelerates for a long period since old duplication for trees with many duplicates. | Performs well for trees with few duplications (proportion 0.2) but lacks power in most cases for trees with proportion of duplication events of 0.5, and of 0.8. |
| Rates of sequence and trait evolution | Rates of sequence evolution, and traits evolutionary rates of events | Fails to show the actual trend of functional evolution under a null, and under certain ortholog conjecture scenario | Recovers the actual trends in all hypothetical conditions |
| Asymmetric trait jump model | Branch of jump and random change in trait between 0-1 | Performs well in all hypothetical scenarios | Recovers the actual signals in all hypothetical conditions |

Unlike under the previous model, the proportion of duplication events in the trees has no impact on the capacity to recover trends (Fig 4, S2-S4 Figs). An important difference on the other hand is that the null is not always properly recovered. When the trait evolutionary rate is not impacted by duplication, but the sequence evolutionary rate is (scenario 2 of Table 1), the PIC approach wrongly rejects the null for trait evolution.

**Fig 4:**
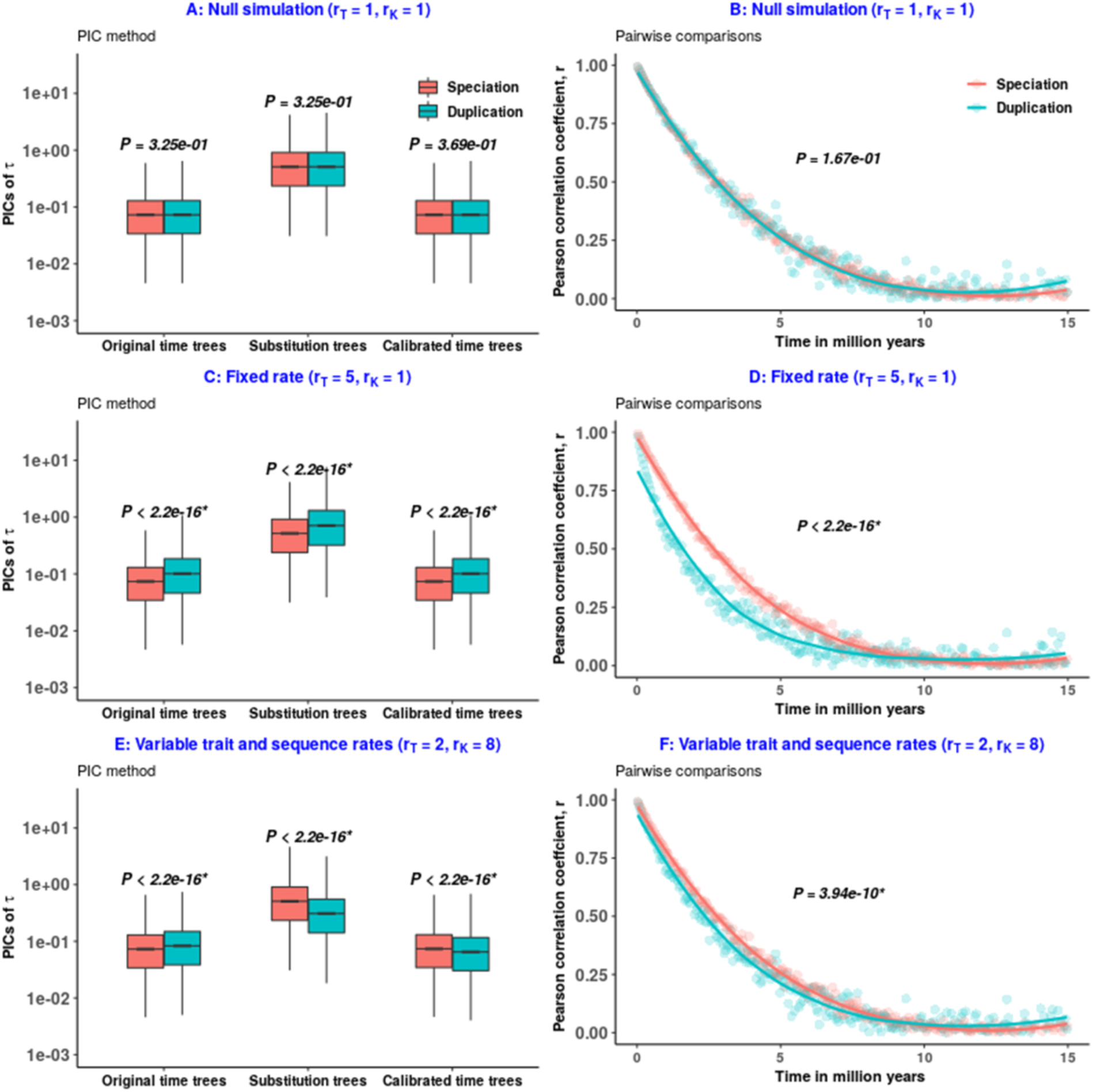
Comparisons of approaches in 3 different scenarios of Table 1. PICs: Phylogenetic Independent Contrasts. We used an asymmetric model, where after duplication one of the two duplicates experiences acceleration. The y-axis is on a log_10_ scale for PICs. A quadratic fit polynomial is used for our improved pairwise comparisons. *: Bonferroni-corrected significant *P* values. A-B. Both the approaches correctly accept the null under a uniform null evolution of sequences and trait. C-D. An ideal ortholog conjecture scenario, i.e. scenario 3 of Table 1; both approaches correctly reject the null in favor of faster evolution after duplication. E-F. The ortholog conjecture scenario 4c of Table 1, where evolutionary rates of sequences accelerate more than those of traits following duplication. E. The PIC correctly rejects the null in favor of faster evolution after duplication when using the true time tree (left); with a tree derived from sequence evolution (middle and right) the PIC rejects the null but incorrectly in favor of faster evolution after speciation. F. Improved pairwise comparisons correctly rejects the null in favor of faster evolution after duplication.

This is due to time tree calibration using sequence evolution as a reference. Indeed the PIC does not reject the null when the original time tree is used (A in S2, C in S3-S6 Figs). However, the higher sequence evolutionary rates after duplication lead to higher expected variance, and thus the calibrated time trees produce lower contrasts for duplication than for speciations, with a significant difference in PIC (A in S2, C in S3-S6 Figs). Moreover, if both the trait and the sequence evolutionary rates accelerate after duplication events, but the sequence acceleration is higher (scenario 4c of Table 1), the PIC can again estimate lower contrasts for duplication branches with calibrated time trees (Fig 4E, K in S3-S6 Figs). This can lead to wrongly accepting the null if a one-sided test is used, or rejecting the null in the wrong direction. This bias does not impact the pairwise approach, under any of these scenarios (Fig 4F, B in S2, D and L in S3-S6 Figs). Thus even though the assumption of independent data points is violated, the improved pairwise approach is more robust than the PIC in this type of model, since it does not depend on the branch lengths.

### Model-3: Asymmetric trait jump model

Finally, we investigated our ability to recover gene evolutionary patterns under a jump model rather than changes in continuous rates. The idea is that, similarly to rapid trait jumps when organisms transition into new adaptive zones (Simpson 1944; Duchen *et al*. 2017), paralogs might undergo rapid trait jumps when they appear, e.g. because of a new chromosomal environment in the case of asymmetric duplication. We used a Brownian motion model with asymmetric jumps in trait (Fig 5). Jumps can follow either speciation or duplication events, and different gene evolution models correspond to different probabilities of jumps after the two types of events. We used a simple model where there can only be one or zero jump per branch. Speciation and duplication branches are randomly chosen, and one of their daughter branches (randomly selected) experiences a rapid jump in trait, then returns to progressive trait evolution along the branch; there is only one progressive evolutionary rate. Under the null, equal proportions of speciation and duplication branches (30%) were randomly chosen. Reassuringly, the null was rejected neither by PIC (A in S7 Fig) nor by the pairwise approach (B in S7 Fig) for the standard pure simulation models. For the ortholog conjecture, we simulated more asymmetric jumps following duplications than speciations. More jumps in trait should introduce more divergence in trait, and this should be detected as support for the ortholog conjecture. Indeed, both approaches did support the ortholog conjecture in this case (C and D in S7 Fig). This was robust to trees with different proportions of duplication events, but not to simulation of trees with empirical data parameters (S8 and S9 Figs). The performance of the PIC is strongly biased by branch lengths from empirical data under this model (A in S10-S11 Figs and C in S10 Fig). The improved pairwise approach is more robust than the PIC in this case, although it lacks power (S10-S11 Figs).

**Fig 5:**
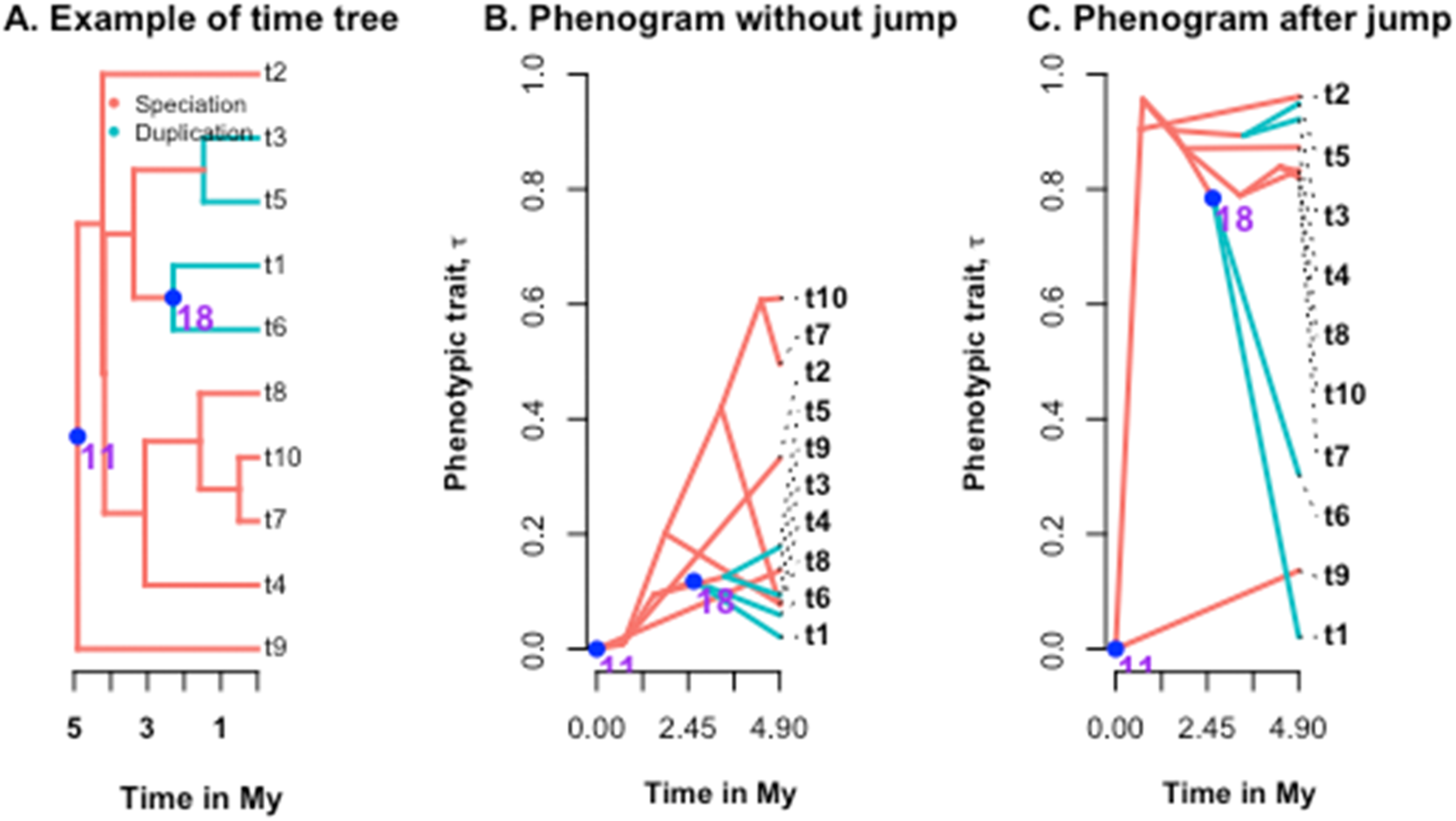
Principle of an asymmetric trait jump model. A. We used a tree with 10 tips (t1-t10) with a proportion of duplications of 0.2 for illustration. This means that the tree has 7 speciation and 2 duplication nodes. To represent an ortholog conjecture scenario, we considered that 20% of the speciation nodes, and 50% of the duplication nodes experienced instantaneous jumps in trait. We chose one of the two branches randomly following the blue colored nodes with numbers, where the trait jump took place. B. The phenogram shows the projection of the phylogenetic tree in a space defined by trait τ of the both events (on the y axis) and evolutionary time (on the x) before the jump in trait. C. The phenogram shows how the trait value changed over the time after experiencing trait jumps. For example, node 11 experienced a trait jump. Hence, the trait value remained unchanged for t9 but alters for the other branch following a jump. This in consequence changed the values of all the following descendants over the time. Hence, the trait value of 18 in C differs from B. For another jump in node 18, t6 changed but not t1.

## Discussion

A good method to detect patterns of gene functional evolution should not reject the null hypothesis when data are simulated under the null, should successfully reject the null when simulated under an alternative hypothesis, and should recover the right direction of pattern difference. Here, our null hypothesis is that gene function evolves independently of duplication, and our alternative is the “ortholog conjecture”, that function evolves faster after duplication. While the most naive form of this ortholog conjecture is one evolutionary rate on duplication branches, and another on speciation branches, we explored three more complex models of trait evolution.

It is worth mentioning that the models under which we have simulated cannot be easily tested on empirical data, since we lack information for the original times trees on which traits have evolved. Moreover, on empirical trees it is impossible to guarantee a true null model including no change in sequence evolutionary rates after duplication. We rather obtain calibrated pseudo-time trees in reality. Such calibrated empirical gene trees cannot always be considered as good proxies of the original time trees, since the branch lengths of such trees are biased, especially for old duplicates, due to the non-availability of reference time points for duplications (Begum & Robinson-Rechavi 2020). When traits are simulated based on such biased calibrated time trees, the performances of both approaches deteriorate by recovering opposite pseudo-signals in the null condition (Begum & Robinson-Rechavi 2020; Dunn *et al*. 2018). Thus they may not be sufficient to trace expected patterns of trait evolution in many realistic scenarios. Even with trees simulated using parameter values from such calibrated time trees, failure to recover the expected trends with sufficient power is common. This is due to the use of biased phylogenies for parameter estimation for simulations. On the contrary, pure simulation models are devoid of such calibration bias, and thus are suitable to investigate the behavior of methods in theoretical scenarios where testing with empirical data seems impossible, including the pure null. If we could obtain calibrated unbiased empirical phylogenies, the trends of functional evolution should be consistent with these pure simulation models for the testable cases, since the pure simulated trees with different proportions of duplications allow to understand the patterns of functional evolution irrespective of the positions of events in trees. We will thus focus on the result from the pure simulated trees.

First, while the jump model is very different from the most naive ortholog conjecture model, both methods tested (pairwise and PIC) performed well under it unless affected by biased phylogenetic patterns (S7-S11 Figs). Thus, testing the ortholog conjecture seems robust to change being sudden or gradual. Second, when gradual change extends in time to both speciation and duplication branches, methods lose power to detect the correct pattern. This loss increases with the duration of acceleration and with the proportion of branches affected. This time dependent trait acceleration model represents a reasonable assumption of gene evolution, as there is no strong reason to expect paralogs to stop evolving like paralogs because there was a speciation. Indeed duplication affects each gene’s direct environment, whereas speciation can have at most a very indirect effect. The PIC was more powerful than the pairwise approach under this model, on pure simulated trees (S2 Table). Thirdly and most worryingly, when both sequence and trait evolutionary rates are affected by duplication, PIC can both incorrectly reject the null under null simulations of trait evolution, and report the opposite direction of change under ortholog conjecture simulations (Fig 4E, A in S2, C and K in S3-S6 Figs).

While all of the simulation models used are quite simple, they represent realistic scenarios of gene function evolution. The mechanisms of trait divergence following gene duplication are still poorly understood. For example, we do not have direct evidence for an asymmetric trait jump model or for a time dependent trait acceleration model. The purpose of our simulation study was to identify scenarios where the PIC and the pairwise comparisons may produce distinct signals of functional evolution between orthologs and paralogs. Although our simple simulations used the same mechanism of trait evolution for all the duplicates of a tree, a mix of different models for duplicates in different parts or at different times in a same gene tree cannot be excluded.

Pairwise comparisons of traits between genes has been criticized in studying functional evolution (Dunn *et al*. 2018). Behavior under a null model can help to assess the presence of a systematic bias in the approach. In all different models, and with different sets of trees (S1 Table), we found that our improved pairwise comparisons gave the expected results under the null. The simple pairwise comparison of Kryuchkova-Mostacci and Robinson-Rechavi (2016) also produced the expected signal under the null (S2 Table), although it lacked power under the alternative model. The largest difference between pairwise and PIC was the failure of PIC when both rates of sequence and of trait are affected by duplication (Fig 4, S2-S6 Figs). This is worrying since acceleration of sequence evolutionary rates is common following gene duplications (Gu *et al*. 2005; Scannell & Wolfe 2008; Panchin *et al*. 2010; Assis & Bachtrog 2013; Pegueroles *et al*. 2013; Kryuchkova-Mostacci & Robinson-Rechavi 2016; Jiang & Assis 2017; Lafond *et al*. 2018). Thus the scenario of these simulations is expected to be widespread in real data. In this case, higher sequence evolutionary rates for duplicates lead to higher expected variances, and thus produce lower PIC for duplication events than speciation events. These lower PIC are in contrast to the null expectation of no difference.

Our simulations also show that in the most probable ortholog conjecture scenario, where both the trait and the sequence evolutionary rates accelerate after duplication events (Gu *et al*. 2005), but the ratio of sequence to the trait acceleration is higher (scenario 4c of Table 1), the PIC method fails to detect the signal of ortholog conjecture (Fig 4E, K in S3-S6 Figs). In this case, improved pairwise comparisons seem to be a better choice due to its branch lengths independence.

The only model for which the ratio of duplication to speciation events impacts the results is with time-dependent trait acceleration. When there are many duplications, and the duplication-induced acceleration lasts for 70% to 90% of old duplication age, most speciation branches also evolve under the influence of trait acceleration like duplicates. Hence, both the approaches can fail to recover the signal of the ortholog conjecture in a real ortholog conjecture scenario (S2 Table). Under intermediate scenarios, PIC allows to better recover the signal of the ortholog conjecture than pairwise comparisons. However, such a pattern is rare in the empirical parameter space we used, since they have low proportions of duplications, and both the approaches performed well under it (S2 Table).

Overall the main message of this study is that our ignorance of gene function evolution makes it difficult to choose one method as the gold standard. Each method has its own limitations, and we call for methodological pluralism, at least in the present state of our knowledge. Relying on a single approach to interpret result can be problematic in many cases of comparative functional genomics study involving complex gene family evolution. These results apply in principle not only to the ortholog conjecture, but to any other cases of gene trait evolution (e.g., horizontal gene transfer), where evolution might be gradual or by jumps, affect a more or less large proportion of branches of the gene tree, and be confounded with changes in sequence evolutionary rates. In conclusion, functional phylogenomics is complex and results supported by only one method should be treated with caution.

## Acknowledgements

We sincerely thank Julien Wollbrett, Elsa Guillot, Sara Fonseca Costa, and all the members of the Robinson-Rechavi and Salamin groups for their help and useful discussions. Parts of the computations were performed at the Vital-IT (http://www.vital-it.ch) Center for high-performance computing of the SIB Swiss Institute of Bioinformatics, as well as using the Wally cluster of the University of Lausanne.

## Author contributions

TB, MLSS, and MRR conceived the ideas; TB and MLSS designed the original methodology; TB and MRR refined the methodology; TB and MLSS wrote the code for analysis with input from MRR; TB analyzed the data; TB and MRR led the writing of the manuscript. All authors contributed critically to the drafts and gave final approval for publication.

## Supporting Information

### Supplementary method details

#### Model-1: Time dependent trait acceleration model

In this model, we used the age of the oldest duplication events in a tree to specify the time of acceleration following gene duplication, since the maximum age varies for each tree. For modeling of “Time dependent trait acceleration”, we wrote our own functions to paint branches of a tree with different colors, so that the trait accelerates until different specified durations of acceleration following duplication in a probable ortholog conjecture scenario. This means that such duplication-caused acceleration will eventually accelerate the evolutionary rates of the successive speciation or NA or duplication events falling within that time period. Trait evolutionary rates of events will revert back to its original rates after that time period. In this model, sequence evolutionary rates do not play any role, and thus do not differ between events (i.e., Y= 1 in S1 Table).

For both types of simulated trees of S1 Table, we simulated the trait, *τ* on each original time tree using a BM. The same trait values were used for the corresponding substitution rate trees, and for the pseudo time calibrated trees for further analyses with phylogenetic independent contrasts method. Since pairwise comparisons do not rely on branch lengths, such comparisons are always based on original time trees.

To test this model, we used a 5 times higher trait evolutionary rate over a specific period of time following gene duplications compared to the speciation events (i.e., σ^2^_duplication_= σ^2^_speciation_* 5) in an ortholog conjecture scenario for the pure simulated trees of S1 Table. The trait evolutionary rates, σ^2^, do not vary between events under the null hypothesis.

To strengthen our conclusion, we arbitrarily used distinct higher trait acceleration rates (i.e., σ^2^_duplication_= σ^2^_speciation_* 1.2, σ^2^_duplication_= σ^2^_speciation_ *2, and σ^2^_duplication_= σ^2^_speciation_ *5, respectively) following duplication over a specific period of time for 6953 trees of S1 Table.

#### Model-2: Rates of sequence and trait evolution model

For modeling of “Rates of sequence and trait evolution” using both types of simulated trees of S1 Table, we randomly painted one of the two branches of the simulated original time trees following gene duplication (Fig 1A). This is to generate asymmetric substitution rates tree (Fig 1B), where the sequence evolutionary rates (K) accelerate only for the painted branches of a chronogram due to duplication-caused acceleration in an ortholog conjecture scenario. Speciation clade ages (as described in methods) were used to generate pseudo time calibrated trees (Fig 1C). We simulated trait on each original time tree using a BM. The same trait values were used for the corresponding substitution rate trees, and for the pseudo time calibrated trees for further analyses with phylogenetic method. In case of the improved pairwise comparisons, we used simulated original time trees to compute the difference in correlation coefficients r between events.

To avoid any bias in our inference, we used the same rates for sequence and trait evolutionary rates in a null scenario (S1 Table). To test this model, we used different ranges of σ^2^, and K values in an ortholog conjecture scenario.

#### Model-3: Asymmetric trait jump model

We used the same trait evolutionary rates for all the events of a tree for modeling of “Asymmetric trait jump” (i.e., σ^2^_duplication =_ σ^2^_speciation_ = *σ^2^*_NA_). In this model, sequence evolutionary rates do not differ between events (i.e., Y= 1 in S1 Table). The *FastBM* function (Revell 2012) was used to simulate the trait τ on each original time tree.

Based on the proportions, speciation and duplication branches of each tree are randomly chosen, and one of their daughter branches (randomly selected) experiences a rapid jump in trait with a range between 0 to 1, then returns to progressive trait evolution along successive branches towards tips with the new ancestral trait values. Consequently, the trait values at tips following those daughter branches alters. We used these modified trait values to contrast the phylogenetic methods and the improved pairwise comparisons.

To test this model using the pure simulated trees of S1 Table, we used equal proportions of speciation and duplication branches (30% *vs.* 30%) as a null, in contrasts to more asymmetric trait jumps following duplication than speciations (50% *vs.* 20%) in an ortholog conjecture scenario. To verify the results for trees simulated with empirical data parameters, we additionally considered 50% *vs.* 50% trait jump as a null, and 80% *vs.* 20% jumps in trait as an ortholog conjecture scenario.

---

**S1 Table:** Information on different tree sets, and their relevant parameters used in this study.

| Dataset | Number of original time trees | Number of calibrated time trees | Proportions of node events | Events annotation type | Type of trait evolutionary rates, $\sigma^2$, for each tree | Trait evolutionary rates, $\sigma^2$, used in different test models | Sequence evolutionary rates, K, used in different test models |
| --- | --- | --- | --- | --- | --- | --- | --- |
| **Pure simulation model**<br>(Speciation rates = 0.4, extinction rates = 0.1) | 10000 | 7711 | $P_{\text{dup}} = 0.2,$<br>$P_{\text{spe}} = 0.8,$<br>$P_{\text{NA}} = 0$ | Arbitrary | Randomly fixed<br>($\sigma^2_{\text{speciation}} = 0.02$) | $\sigma^2_{\text{duplication}} = X * \sigma^2_{\text{speciation}}$ | $K_{\text{speciation}} = \sigma^2_{\text{speciation}} = 0.02,$<br>$K_{\text{duplication}} = Y * K_{\text{speciation}}$ |
| | 10000 | 6752 | $P_{\text{dup}} = 0.5,$<br>$P_{\text{spe}} = 0.5,$<br>$P_{\text{NA}} = 0$ | | Randomly fixed<br>($\sigma^2_{\text{speciation}} = 0.02$) | | |
| | 10000 | 8251 | $P_{\text{dup}} = 0.8,$<br>$P_{\text{spe}} = 0.2,$<br>$P_{\text{NA}} = 0$ | | Randomly fixed<br>($\sigma^2_{\text{speciation}} = 0.02$) | | |
| **Simulations based on empirical data parameters**<br>(Speciation rates = 0.013, extinction rates = 0.007) | 6953 | 6953 | $P_{\text{dup}} = 0.1,$<br>$P_{\text{spe}} = 0.7,$<br>$P_{\text{NA}} = 0.2$ | Arbitrary | Estimated using empirical $\tau$ of tips<br>($\sigma^2_{\text{speciation}} = \sigma^2_{\text{calibrated empirical tree}}$) | $\sigma^2_{\text{duplication}} = X * \sigma^2_{\text{speciation}},$<br>$\sigma^2_{\text{NA}} = \sigma^2_{\text{speciation}}$ | $K_{\text{speciation}} = \sigma^2_{\text{speciation}},$<br>$K_{\text{duplication}} = Y * K_{\text{speciation}},$<br>$K_{\text{NA}} = K_{\text{speciation}}$ |

*Note* - $P_{\text{dup}}$ : Proportion of duplications; $P_{\text{spe}}$ : Proportion of speciations; $P_{\text{NA}}$ : Proportion of NA events; X: Multipliers of $\sigma^2$ , Y: Multipliers of K.
X= 1 and Y= 1 in a null condition of any of the 3 test models. Y=1 for “time dependent trait acceleration model”, and for “asymmetric trait jump
model”.

**S2 Table:** Performances of the PIC and pairwise (PW) approaches for time dependent trait acceleration model based on pure simulations.

| | ***P* values from proportion of duplications = 0.2, where**<br>$\sigma^2_{\text{duplication}} = 5 * \sigma^2_{\text{speciation}},$<br>$\sigma^2_{\text{speciation}} = 0.02$ | | | ***P* values from proportion of duplications = 0.5, where**<br>$\sigma^2_{\text{duplication}} = 5 * \sigma^2_{\text{speciation}},$<br>$\sigma^2_{\text{speciation}} = 0.02$ | | | ***P* values from proportion of duplications = 0.8, where**<br>$\sigma^2_{\text{duplication}} = 5 * \sigma^2_{\text{speciation}},$<br>$\sigma^2_{\text{speciation}} = 0.02$ | | |
| --- | --- | --- | --- | --- | --- | --- | --- | --- | --- |
| Approach | PIC | PW | PW_KMR<br>R | PIC | PW | PW_KMR<br>R | PIC | PW | PW_KM<br>RR |
| **Null hypothesis** | 3.69e<sup>-01</sup> | 1.54e<sup>-01</sup> | 6.12e<sup>-01</sup> | 6.78e<sup>-01</sup> | 3.50e<sup>-01</sup> | 7.44e<sup>-01</sup> | 3.71e<sup>-01</sup> | 2.15e<sup>-01</sup> | 4.73e<sup>-01</sup> |
| **Trait acceleration for 50% time of old duplication age** | < 2.2e<sup>-16*</sup> | < 2.2e<sup>-16*</sup> | 1.11e<sup>-4*</sup> | < 2.2e<sup>-16*</sup> | 3.74e<sup>-05*</sup> | 2.46e<sup>-01</sup> | 4.03e<sup>-12*</sup> | 7.33e<sup>-01</sup> | 6.05e<sup>-01</sup> |
| **Trait acceleration for 60% time of old duplication age** | < 2.2e<sup>-16*</sup> | < 2.2e<sup>-16*</sup> | 4.62e<sup>-4</sup> | < 2.2e<sup>-16*</sup> | 3.22e<sup>-03</sup> | 4.20e<sup>-01</sup> | 1.06e<sup>-04*</sup> | 9.44e<sup>-01</sup> | 6.92e<sup>-01</sup> |
| **Trait acceleration for 70% time of old duplication age** | < 2.2e<sup>-16*</sup> | < 2.2e<sup>-16*</sup> | 2.04e<sup>-3</sup> | < 2.2e<sup>-16*</sup> | 2.22e<sup>-01</sup> | 5.89e<sup>-01</sup> | 1.17e<sup>-01</sup> | 9.28e<sup>-01</sup> | 7.49e<sup>-01</sup> |
| **Trait acceleration**<br>**for 80% time of**<br>**old duplication**<br>**age** | $< 2.2\text{e}^{-16*}$ | $< 2.2\text{e}^{-16*}$ | $6.00\text{e}^{-3}$ | $< 2.2\text{e}^{-16*}$ | $4.1\text{e}^{-01}$ | $7.1\text{e}^{-01}$ | $9.06\text{e}^{-01}$ | $8.62\text{e}^{-01}$ | $7.8\text{e}^{-01}$ |
| **Trait acceleration**<br>**for 90% time of**<br>**old duplication**<br>**age** | $< 2.2\text{e}^{-16*}$ | $< 2.2\text{e}^{-16*}$ | $1.89\text{e}^{-2}$ | $2.91\text{e}^{-11*}$ | $5.69\text{e}^{-01}$ | $7.97\text{e}^{-01}$ | $6.39\text{e}^{-01}$ | $8.17\text{e}^{-01}$ | $8.00\text{e}^{-01}$ |

*Note:* PIC: Phylogenetic Independent Contrasts method. PW: Improved pairwise comparisons. PW\_KMRR: pairwise comparisons as used by Kryuchkova-Mostacci and Robinson-Rechavi (2016). *P* values of PIC method are shown for calibrated time trees. Due to independence of branch lengths, *P* values of pairwise comparisons are estimated based on original time trees. \*: Bonferroni-corrected significant *P* values.

**S3 Table:**
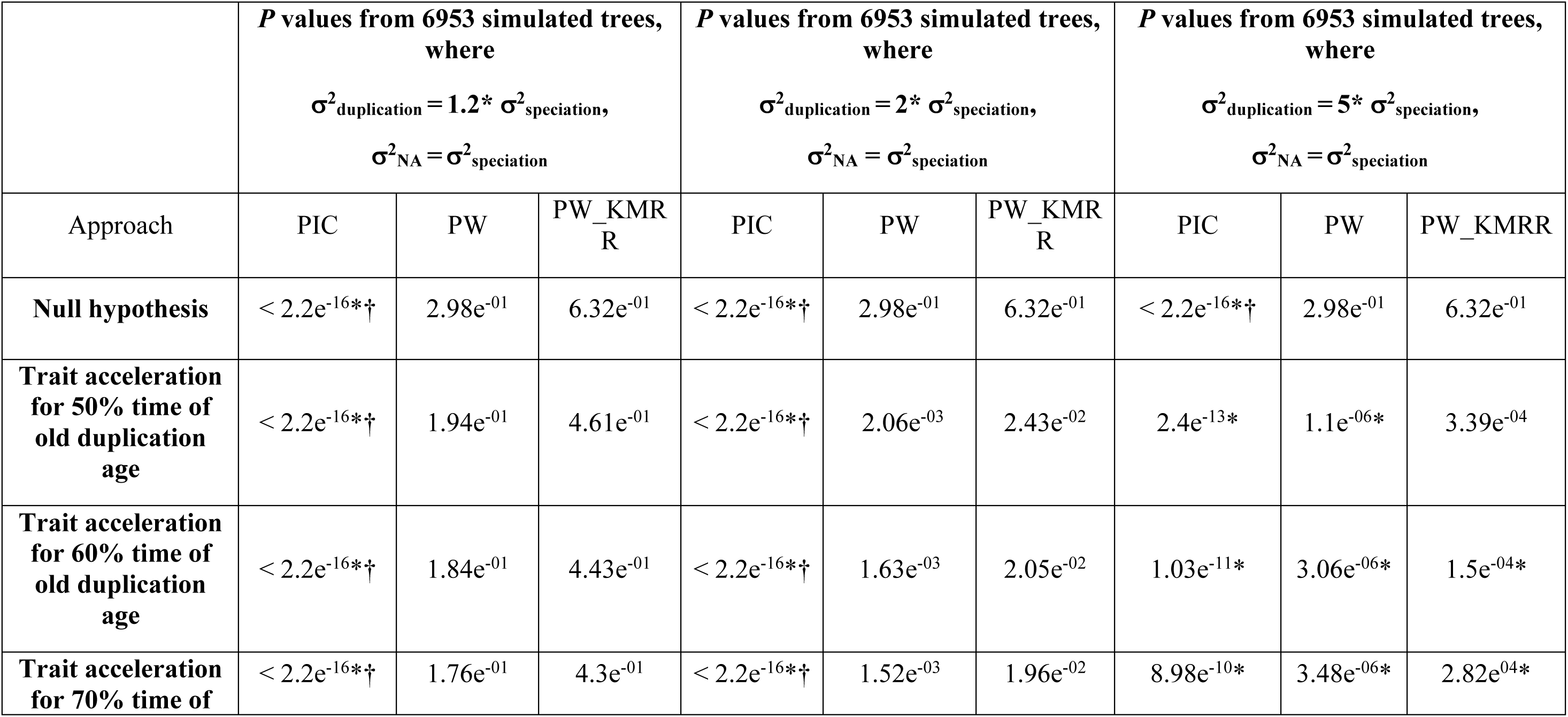
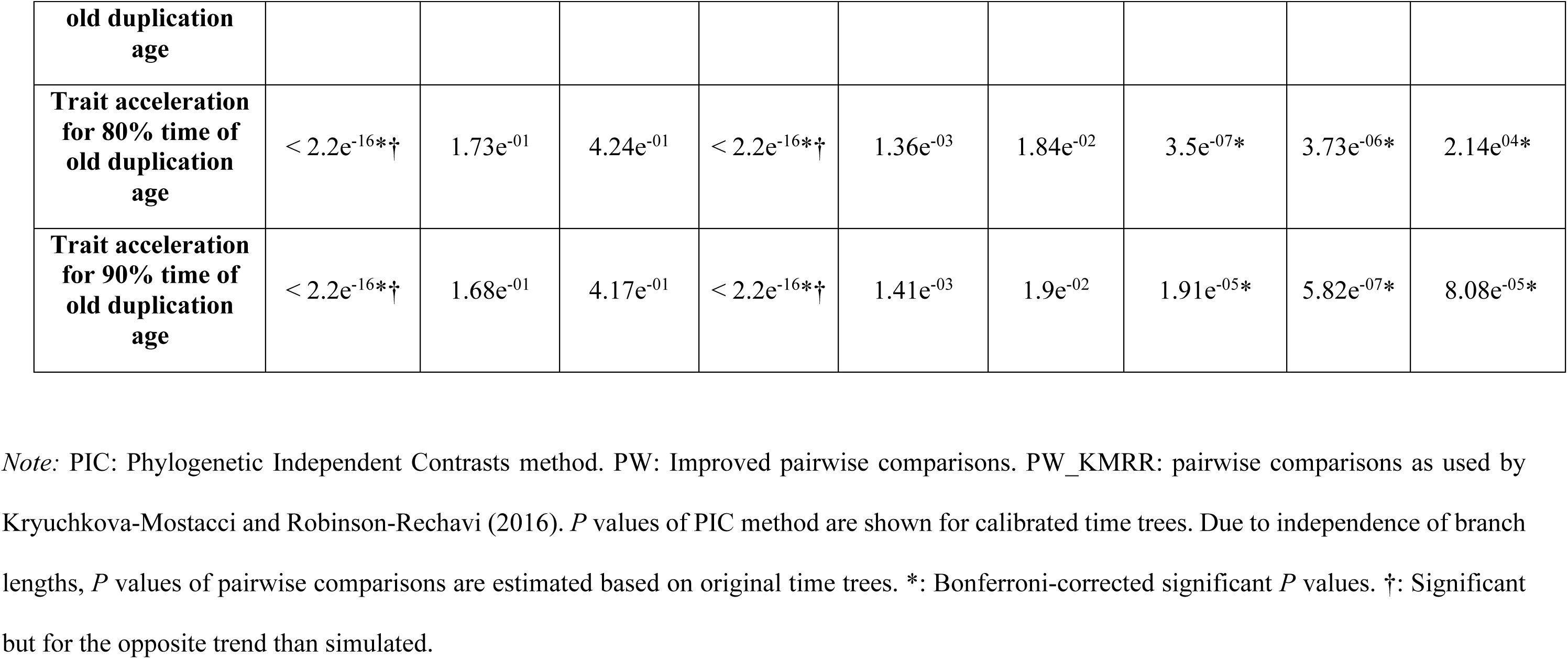
Performances of the PIC and pairwise (PW) approaches for time dependent trait acceleration model based on simulations with empirical data parameters.

## Supplementary figure legends

**S1 Fig**: We used same simulated tree with 100 tips to illustrate 3 different sets (A-C). The branch lengths of the trees are in million years, My. For A-C, the proportion of duplications are 0.2, 0.5, and 0.8, respectively. This means that out of 99 internal nodes 20 nodes, 50 nodes, and 80 nodes are duplication events, respectively in A-C. Rest of the nodes are annotated as speciation nodes.

**Fig S2**: Comparisons of approaches for pure simulated trees with proportion of duplications of 0.2. PICs: phylogenetic independent contrasts. The y-axis is on a log10 scale for the phylogenetic method. *: Bonferroni-corrected significant *P* values. We used an asymmetric model, where after duplication one among the two duplicates experience acceleration. A quadratic polynomial is used in our improved pairwise approach. A-B. Scenario 2 of Table 1, A. Due to no variation in trait evolutionary rates following speciation and duplication events, the original time trees recover an expected signal of no difference in contrasts. An excess of sequence evolutionary rates following duplication than speciation produces lower PICs for duplications compared to speciations for the substitution rate trees, and for the calibrated time trees. This leads to a false rejection of the null in a real null condition for the calibrated trees. B. The improved pairwise approach shows the expected trend by showing no difference in correlation coefficients between events in this case. C-F. The ortholog conjecture scenario 4a and 4b of Table 1. When the trait evolutionary rates ratio is higher than (C-D), or equals to (E-F) the sequence evolutionary rates ratio of events, higher changes in trait rates following duplications make both the approaches to recover the expected signals in support of the conjecture.

**Fig S3**: Performances of the PIC *vs.* improved pairwise approaches in all hypothetical conditions of Table 1. These plots are generated for pure simulated trees with proportion of duplication events of 0.5. Results are for the asymmetric model. PICs: Phylogenetic independent contrasts. *: Bonferroni-corrected significant *P* values. The y-axis is on a log10 scale for the phylogenetic method. We used a quadratic fit in our improved pairwise approach.

**Fig S4**: Performances of the PIC *vs.* improved pairwise methods in the 6 scenarios of Table 1. We used pure simulated trees with proportion of duplication events of 0.8. Results are for the asymmetric model. PICs: Phylogenetic independent contrasts. *: Bonferroni-corrected significant *P* values. The y-axis is on a log10 scale for the phylogenetic method. We used a quadratic fit curve in our improved pairwise comparisons.

**Fig S5**: Performances of the PIC *vs.* improved pairwise methods in the 6 scenarios of Table 1. We used 6953 trees simulated using empirical data parameters. Results are for the asymmetric model. PICs: Phylogenetic independent contrasts. *: Bonferroni-corrected significant *P* values. The y-axis is on a log10 scale for the phylogenetic method. We used a cubic fit curve in our improved pairwise comparisons.

**Fig S6**: Performances of the PIC *vs.* improved pairwise methods in the 6 scenarios of Table 1. We used 6953 simulated trees with different parameters settings compared to Fig S5. Both the methods show same trends of results, as observed from pure simulated trees with different proportions of duplications (Fig 4, S2-S4 Figs), except for the pure null condition, where the phylogenetic method fails. Results are for the asymmetric model. PICs: Phylogenetic independent contrasts. *: Bonferroni-corrected significant *P* values. The y-axis is on a log10 scale for the phylogenetic method. We used a cubic fit curve in our improved pairwise comparisons.

**Fig S7**: Comparisons of the approaches for an asymmetric trait jump model. PICs: Phylogenetic independent contrasts. We used here pure simulated trees with proportion of duplications of 0.2. A quadratic fit curve is used for the improved pairwise approach. *: Bonferroni-corrected significant *P* values. For a null uniform model (A-B), equal proportions, i.e. 30% of the speciation and duplication nodes experienced jump in trait. For the ortholog conjecture scenario, we used higher proportions of trait jumps following duplication nodes than speciation nodes, i.e. 50% and 20% in this case (C-D). Both the approaches recovered the expected signals in all the conditions.

**Fig S8**: Performances of the PIC *vs.* improved pairwise comparisons for an asymmetric trait jump model. PICs: Phylogenetic independent contrasts. We here used pure simulated trees with proportion of duplications of 0.5. We used quadratic fit curve in our improved pairwise comparisons. *: Bonferroni-corrected significant *P* values. (A-B) For a null uniform model, 30% of the nodes experienced trait jump following speciation, and duplication events. PICs or correlations of both events are drawn from the same distribution in such case. For the ortholog conjecture simulation (C-D), 20% of the speciation and 50% of the duplication nodes experienced jumps in trait. Both approaches recover expected signals in all cases.

**Fig S9**: PICs vs. improved pairwise comparisons under an asymmetric trait jump model. PICs: Phylogenetic independent contrasts. We here used pure simulated trees with a proportion of duplications of 0.8. *: Bonferroni-corrected significant *P* values. A quadratic fit curve is used for the improved pairwise comparisons. 30% of the speciation and duplication nodes experienced jumps in trait under a null (A-B), while 20% of the speciation and 50% of the duplication nodes experienced jumps under an ortholog conjecture simulation (C-D). Both the approaches recovered expected signals in all cases.

**Fig S10**: PICs vs. improved pairwise comparisons under an asymmetric trait jump model for 6953 simulated trees. PICs: Phylogenetic independent contrasts. *: Bonferroni-corrected significant *P* values. A cubic fit curve is used for the improved pairwise comparisons. 30% of the speciation and duplication nodes experienced jumps in trait under a null (A-B), while 20% of the speciation and 50% of the duplication nodes experienced jumps under an ortholog conjecture simulation (C-D). An improved pairwise approach is more robust compared to the phylogenetic method in this case.

**Fig S11**: PICs vs. improved pairwise comparisons under an asymmetric trait jump model. PICs: Phylogenetic independent contrasts. We here used 6953 trees that were simulated with empirical data parameters *: Bonferroni-corrected significant *P* values. A cubic fit curve is used for the improved pairwise comparisons. 50% of the speciation and duplication nodes experienced jumps in trait under a null (A-B), while 20% of the speciation and 80% of the duplication nodes experienced jumps under an ortholog conjecture simulation (C-D). An improved pairwise comparisons detect the actual trends of functional evolution in all cases, with a better power of phylogenetic contrasts under the ortholog conjecture scenario.

