## supplementary figures for "Performance of a phylogenetic independent contrast method and an improved pairwise comparison under different scenarios of trait evolution after speciation and duplication"

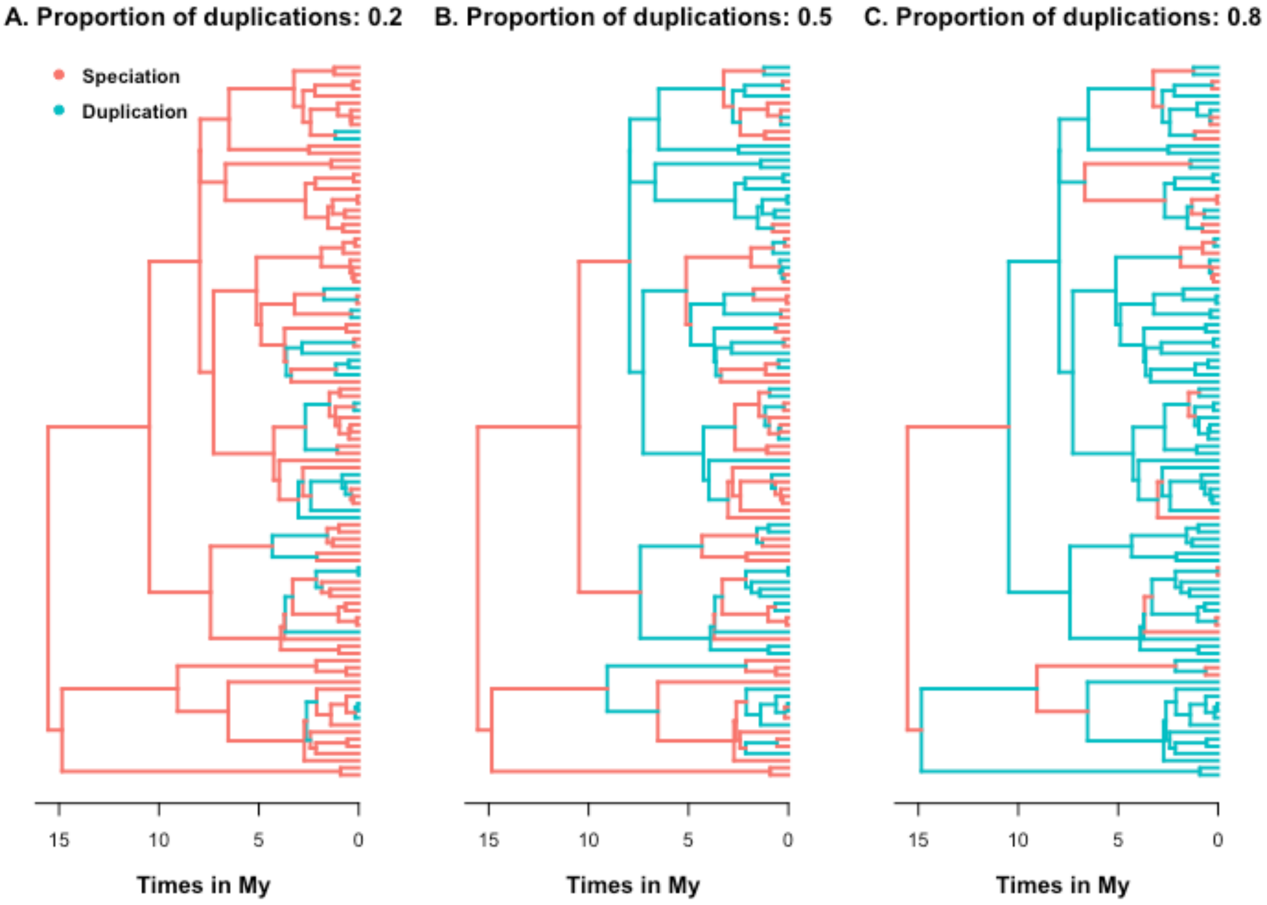

A. Proportion of duplications: 0.2
B. Proportion of duplications: 0.5
C. Proportion of duplications: 0.8
Speciation
Duplication
15
10
5
0
Times in My


**Fig S1**

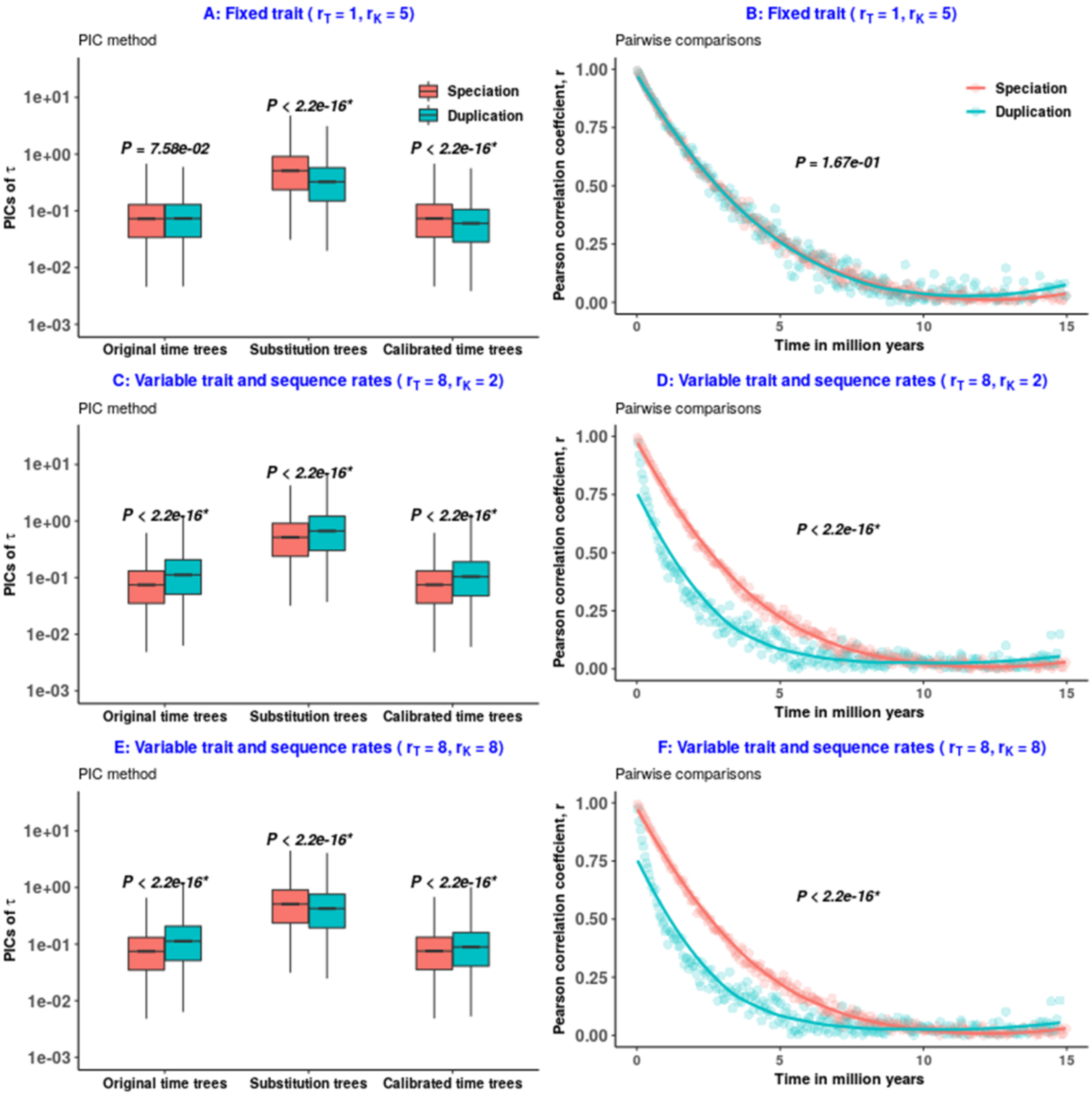
A: Fixed trait ( rT = 1, rK = 5)
PIC method
PICs of τ
P = 7.58e-02
P < 2.2e-16*
P < 2.2e-16*
Speciation
Duplication
Original time trees
Substitution trees
Calibrated time trees
B: Fixed trait ( rT = 1, rK = 5)
Pairwise comparisons
Pearson correlation coeffcient, r
P = 1.67e-01
Time in million years
C: Variable trait and sequence rates ( rT = 8, rK = 2)
P < 2.2e-16*
D: Variable trait and sequence rates ( rT = 8, rK = 2)
E: Variable trait and sequence rates ( rT = 8, rK = 8)
F: Variable trait and sequence rates ( rT = 8, rK = 8)

**Fig S2**

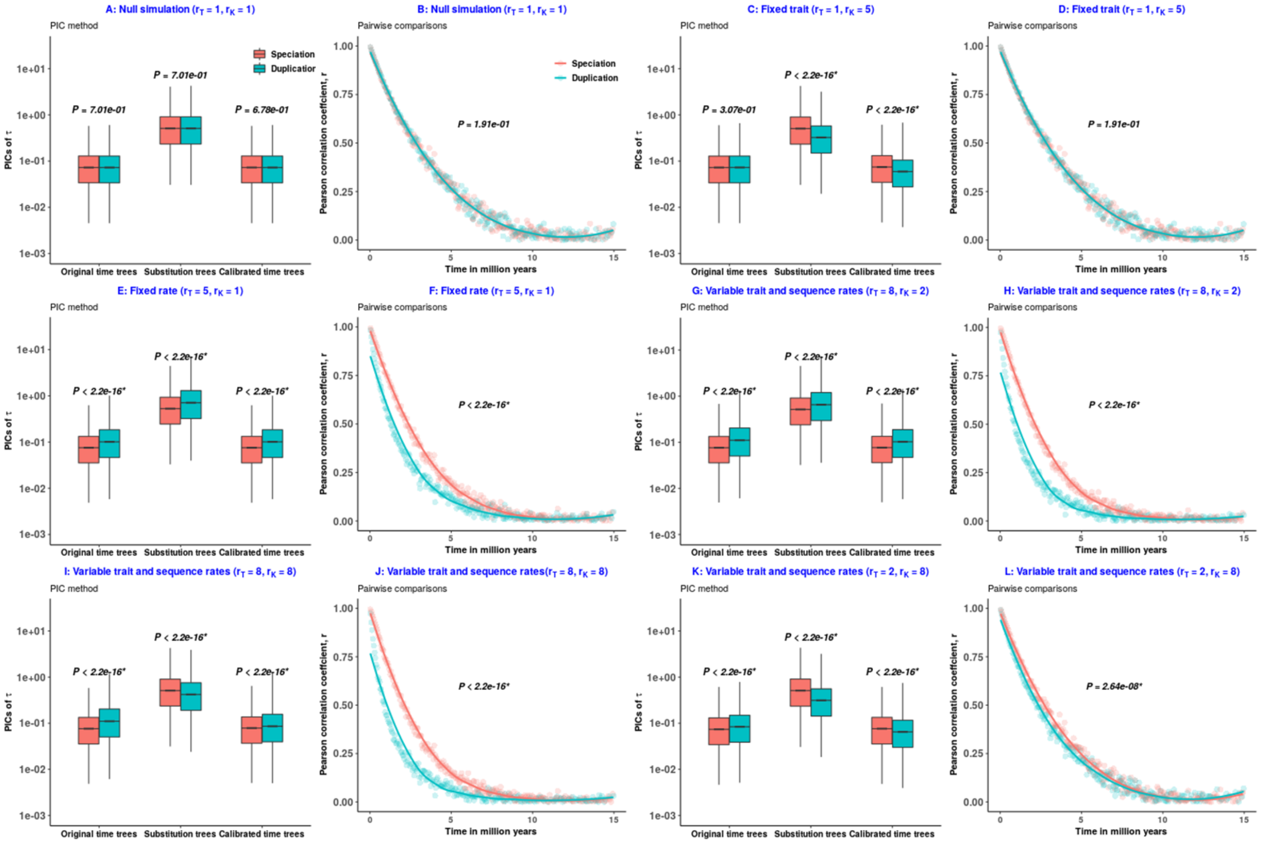

A: Null simulation (rT = 1, rK = 1)
PIC method
Speciation
Duplication
P = 7.01e-01
P = 7.01e-01
P = 6.78e-01
PICs of τ
Original time trees
Substitution trees
Calibrated time trees
B: Null simulation (rT = 1, rK = 1)
Pairwise comparisons
P = 1.91e-01
Pearson correlation coefficient, r
Time in million years
C: Fixed trait (rT = 1, rK = 5)
P = 3.07e-01
P < 2.2e-16*
P < 2.2e-16*
D: Fixed trait (rT = 1, rK = 5)
P = 1.91e-01
E: Fixed rate (rT = 5, rK = 1)
P < 2.2e-16*
F: Fixed rate (rT = 5, rK = 1)
G: Variable trait and sequence rates (rT = 8, rK = 2)
H: Variable trait and sequence rates (rT = 8, rK = 2)
I: Variable trait and sequence rates (rT = 8, rK = 8)
J: Variable trait and sequence rates(rT = 8, rK = 8)
K: Variable trait and sequence rates (rT = 2, rK = 8)
L: Variable trait and sequence rates (rT = 2, rK = 8)
P = 2.64e-08*


Fig S3

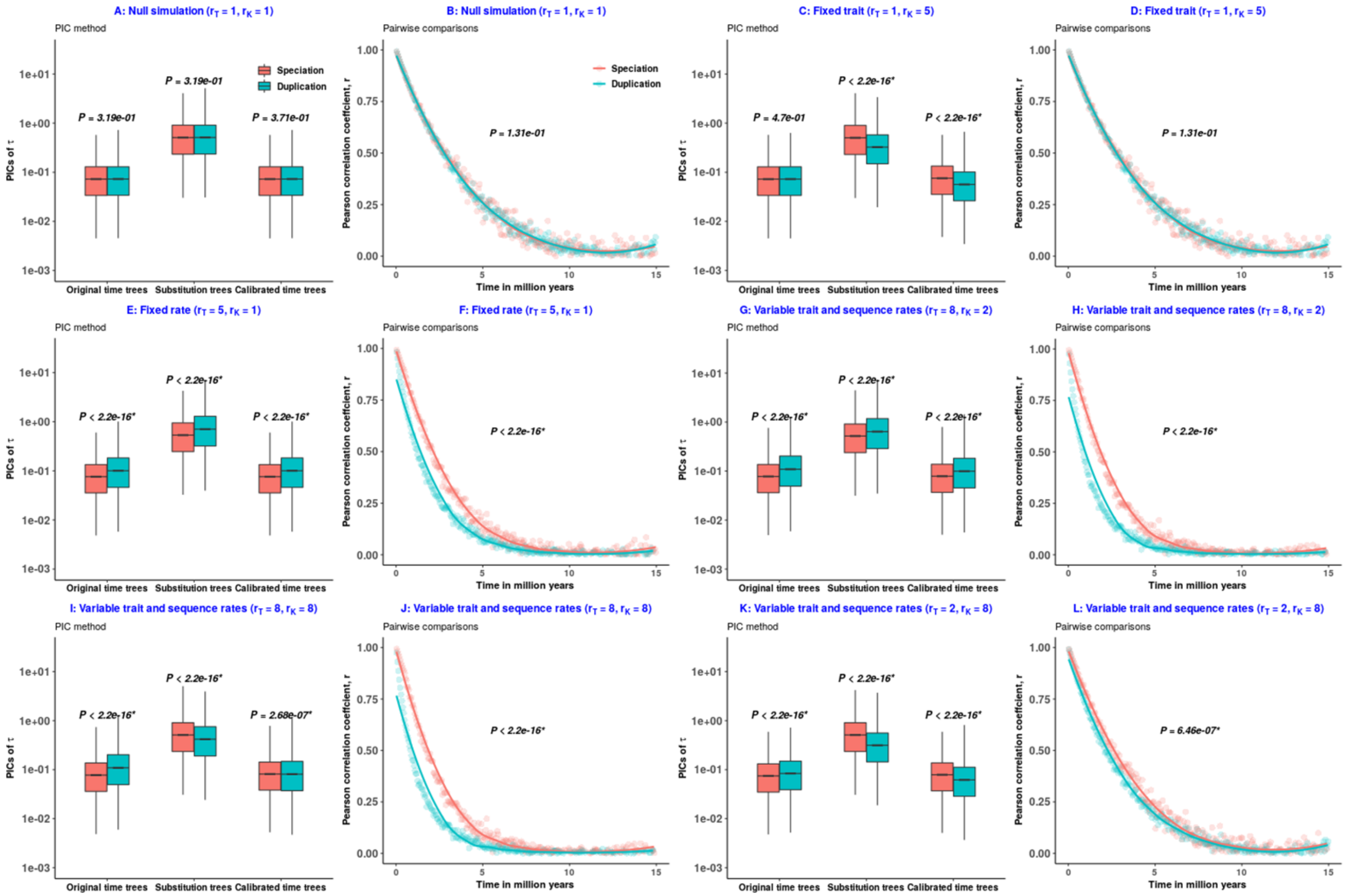

A: Null simulation ($r_T$ = 1, $r_K$ = 1)
PIC method
Speciation
Duplication
P = 3.19e-01
P = 3.19e-01
P = 3.71e-01
PICs of τ
Original time trees
Substitution trees
Calibrated time trees
B: Null simulation ($r_T$ = 1, $r_K$ = 1)
Pairwise comparisons
P = 1.31e-01
Pearson correlation coefficient, r
Time in million years
C: Fixed trait ($r_T$ = 1, $r_K$ = 5)
PIC method
P = 4.7e-01
P < 2.2e-16*
P < 2.2e-16*
D: Fixed trait ($r_T$ = 1, $r_K$ = 5)
Pairwise comparisons
P = 1.31e-01
E: Fixed rate ($r_T$ = 5, $r_K$ = 1)
PIC method
P < 2.2e-16*
P < 2.2e-16*
P < 2.2e-16*
F: Fixed rate ($r_T$ = 5, $r_K$ = 1)
Pairwise comparisons
P < 2.2e-16*
G: Variable trait and sequence rates ($r_T$ = 8, $r_K$ = 2)
PIC method
P < 2.2e-16*
P < 2.2e-16*
P < 2.2e-16*
H: Variable trait and sequence rates ($r_T$ = 8, $r_K$ = 2)
Pairwise comparisons
P < 2.2e-16*
I: Variable trait and sequence rates ($r_T$ = 8, $r_K$ = 8)
PIC method
P < 2.2e-16*
P < 2.2e-16*
P = 2.68e-07*
J: Variable trait and sequence rates ($r_T$ = 8, $r_K$ = 8)
Pairwise comparisons
P < 2.2e-16*
K: Variable trait and sequence rates ($r_T$ = 2, $r_K$ = 8)
PIC method
P < 2.2e-16*
P < 2.2e-16*
P < 2.2e-16*
L: Variable trait and sequence rates ($r_T$ = 2, $r_K$ = 8)
Pairwise comparisons
P = 6.46e-07*


Fig S4

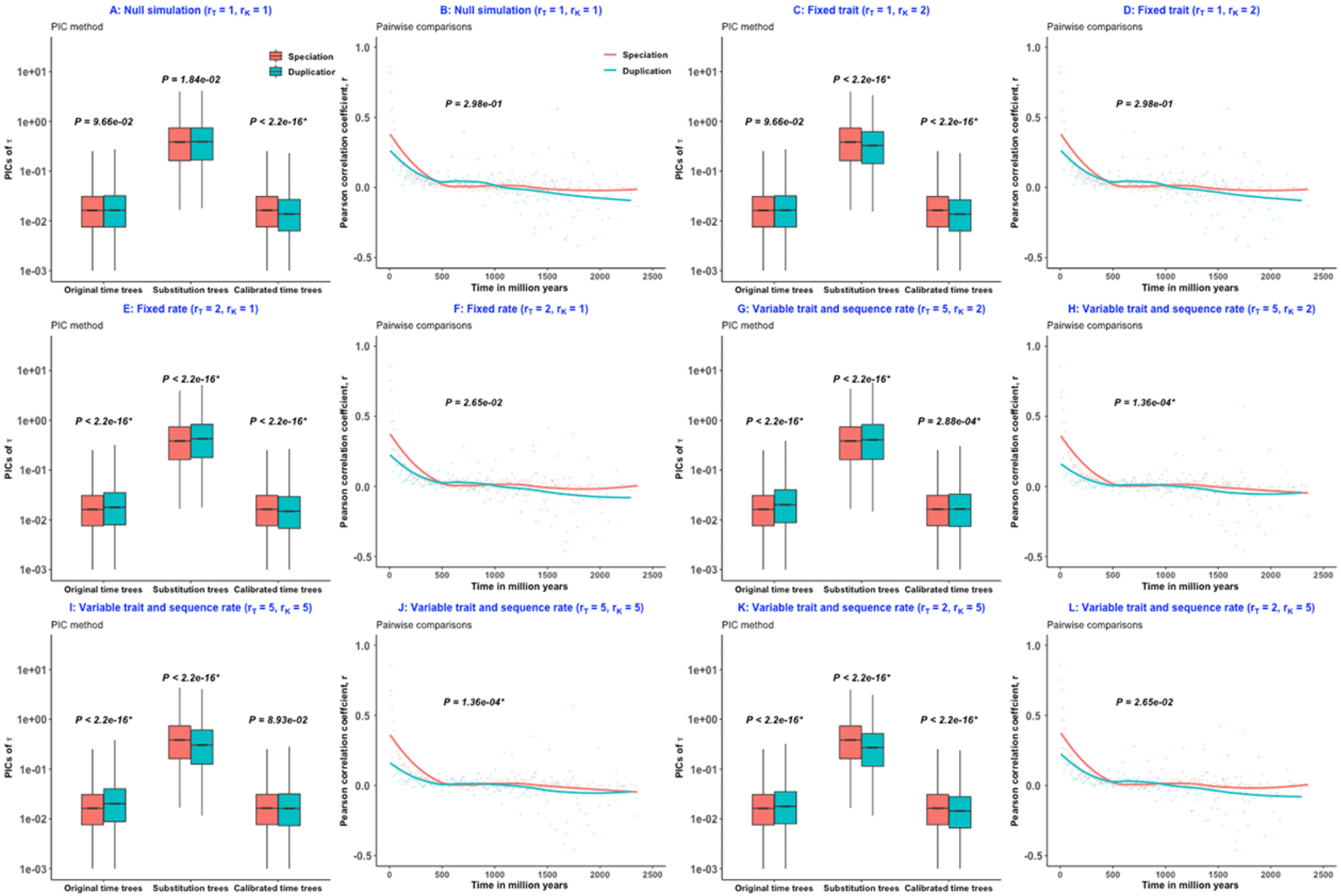

A: Null simulation ($r_T$ = 1, $r_K$ = 1)
B: Null simulation ($r_T$ = 1, $r_K$ = 1)
C: Fixed trait ($r_T$ = 1, $r_K$ = 2)
D: Fixed trait ($r_T$ = 1, $r_K$ = 2)
E: Fixed rate ($r_T$ = 2, $r_K$ = 1)
F: Fixed rate ($r_T$ = 2, $r_K$ = 1)
G: Variable trait and sequence rate ($r_T$ = 5, $r_K$ = 2)
H: Variable trait and sequence rate ($r_T$ = 5, $r_K$ = 2)
I: Variable trait and sequence rate ($r_T$ = 5, $r_K$ = 5)
J: Variable trait and sequence rate ($r_T$ = 5, $r_K$ = 5)
K: Variable trait and sequence rate ($r_T$ = 2, $r_K$ = 5)
L: Variable trait and sequence rate ($r_T$ = 2, $r_K$ = 5)
PIC method
Pairwise comparisons
Speciation
Duplication
PICs of τ
Pearson correlation coeffcient, r
Original time trees
Substitution trees
Calibrated time trees
Time in million years
P = 9.66e-02
P = 1.84e-02
P < 2.2e-16*
P = 2.98e-01
P = 2.65e-02
P = 2.88e-04*
P = 1.36e-04*
P = 8.93e-02


**Fig S5**

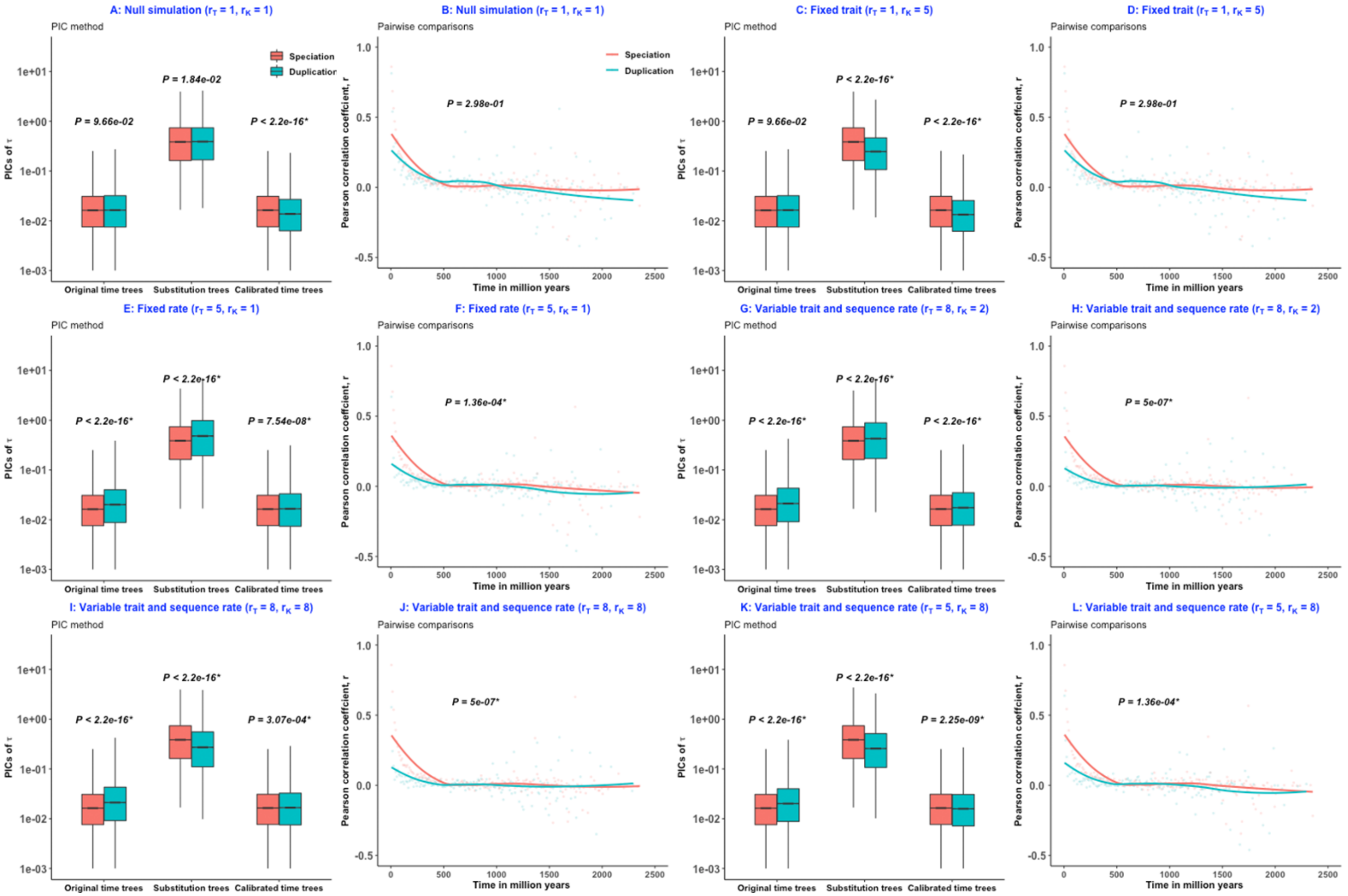
A: Null simulation ($r_T$ = 1, $r_K$ = 1)
PIC method
B: Null simulation ($r_T$ = 1, $r_K$ = 1)
Pairwise comparisons
C: Fixed trait ($r_T$ = 1, $r_K$ = 5)
PIC method
D: Fixed trait ($r_T$ = 1, $r_K$ = 5)
Pairwise comparisons
E: Fixed rate ($r_T$ = 5, $r_K$ = 1)
PIC method
F: Fixed rate ($r_T$ = 5, $r_K$ = 1)
Pairwise comparisons
G: Variable trait and sequence rate ($r_T$ = 8, $r_K$ = 2)
PIC method
H: Variable trait and sequence rate ($r_T$ = 8, $r_K$ = 2)
Pairwise comparisons
I: Variable trait and sequence rate ($r_T$ = 8, $r_K$ = 8)
PIC method
J: Variable trait and sequence rate ($r_T$ = 8, $r_K$ = 8)
Pairwise comparisons
K: Variable trait and sequence rate ($r_T$ = 5, $r_K$ = 8)
PIC method
L: Variable trait and sequence rate ($r_T$ = 5, $r_K$ = 8)
Pairwise comparisons
Speciation
Duplication
PICs of τ
Pearson correlation coefficient, r
Original time trees
Substitution trees
Calibrated time trees
Time in million years
P = 9.66e-02
P = 1.84e-02
P < 2.2e-16*
P = 2.98e-01
P = 7.54e-08*
P = 1.36e-04*
P = 5e-07*
P = 3.07e-04*
P = 2.25e-09*

Fig S6

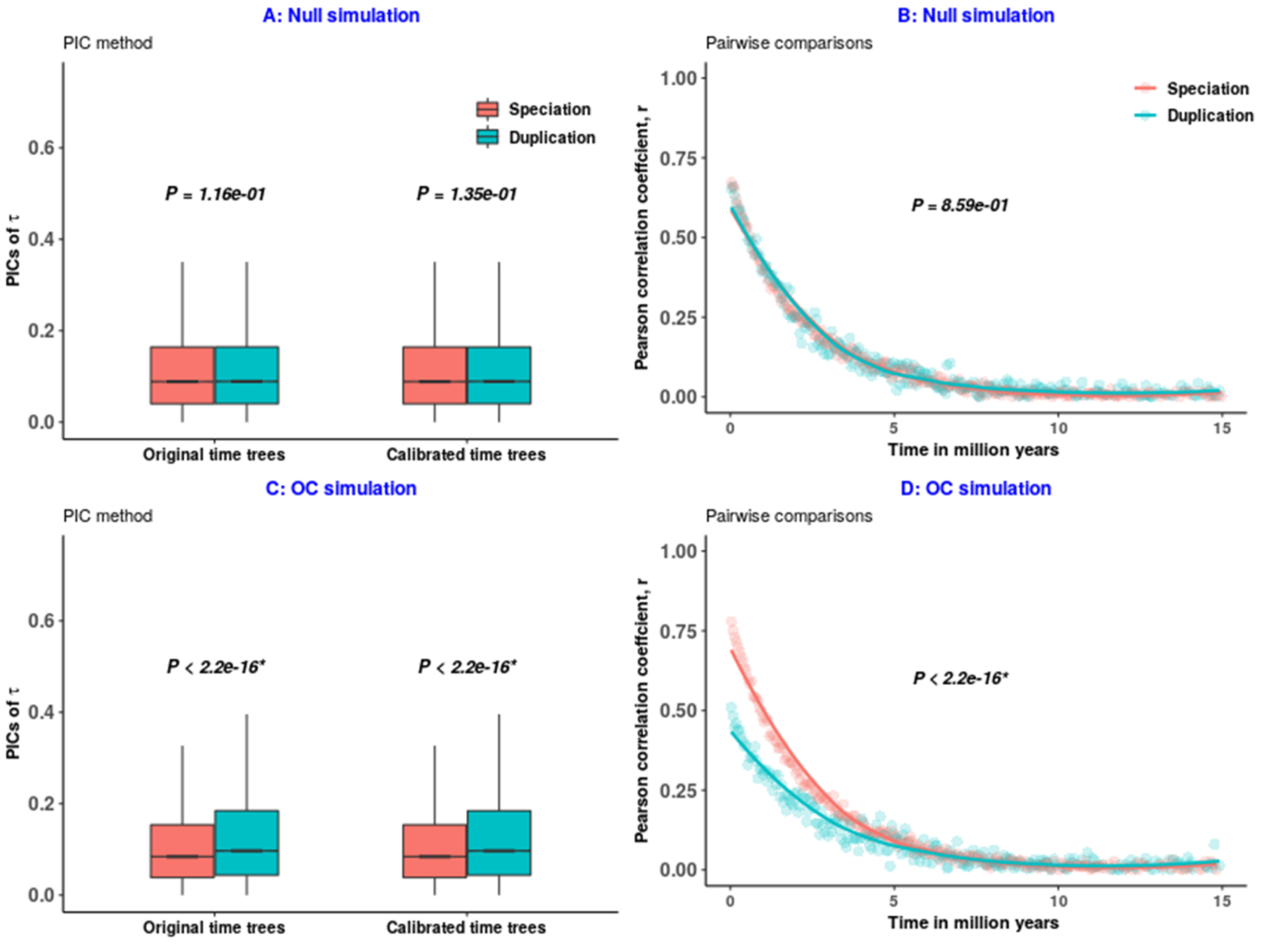
A: Null simulation
PIC method
Speciation
Duplication
P = 1.16e-01
P = 1.35e-01
PICs of τ
0.0
0.2
0.4
0.6
Original time trees
Calibrated time trees
B: Null simulation
Pairwise comparisons
Speciation
Duplication
P = 8.59e-01
Pearson correlation coeffcient, r
1.00
0.75
0.50
0.25
0.00
0
5
10
15
Time in million years
C: OC simulation
PIC method
P < 2.2e-16*
P < 2.2e-16*
PICs of τ
0.0
0.2
0.4
0.6
Original time trees
Calibrated time trees
D: OC simulation
Pairwise comparisons
P < 2.2e-16*
Pearson correlation coeffcient, r
1.00
0.75
0.50
0.25
0.00
0
5
10
15
Time in million years

**Fig S7**

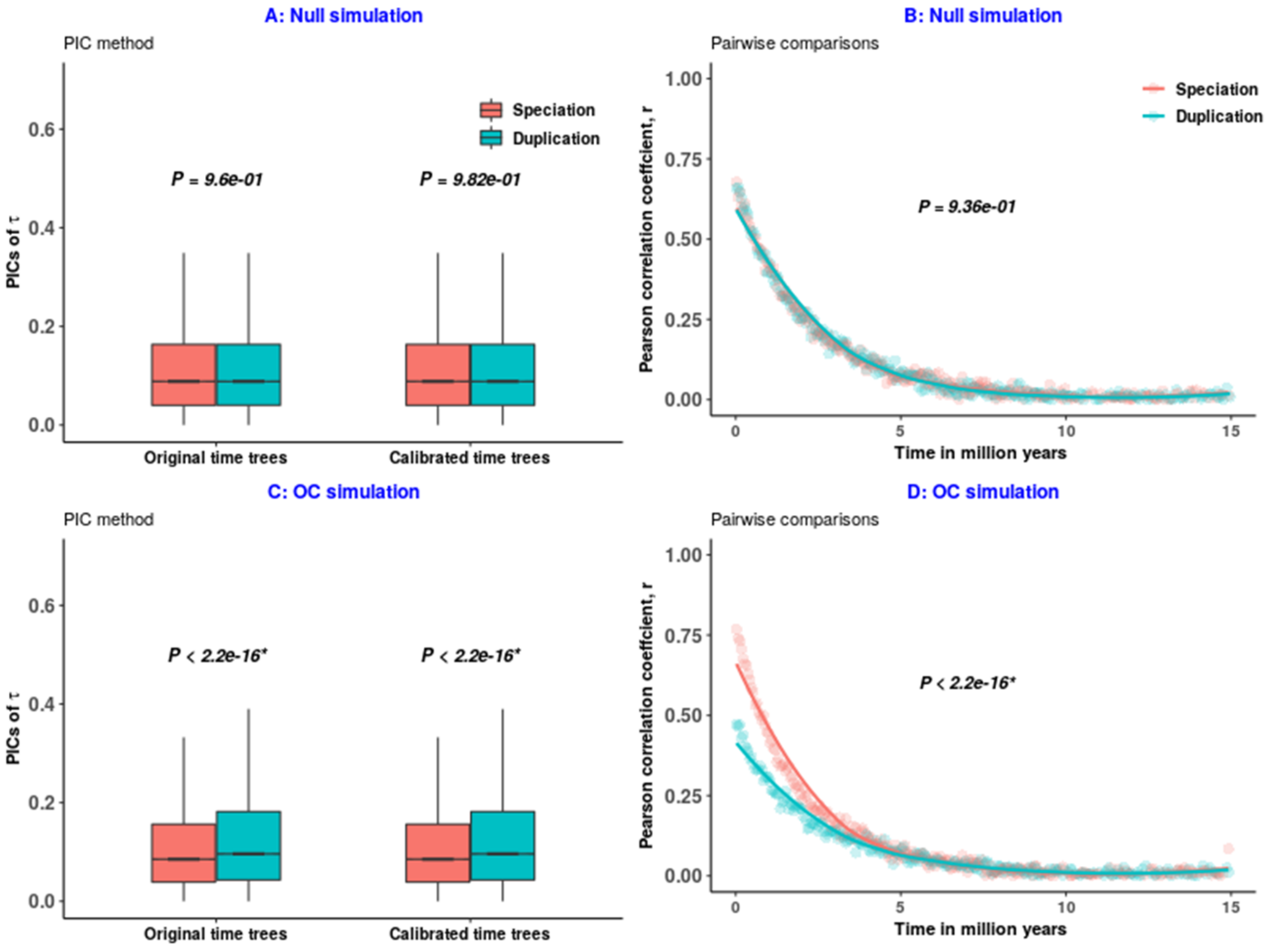
A: Null simulation
PIC method
PICs of τ
0.6
0.4
0.2
0.0
Speciation
Duplication
P = 9.6e-01
P = 9.82e-01
Original time trees
Calibrated time trees
B: Null simulation
Pairwise comparisons
Pearson correlation coeffcient, r
1.00
0.75
0.50
0.25
0.00
Speciation
Duplication
P = 9.36e-01
0
5
10
15
Time in million years
C: OC simulation
PIC method
PICs of τ
P < 2.2e-16*
P < 2.2e-16*
Original time trees
Calibrated time trees
D: OC simulation
Pairwise comparisons
Pearson correlation coeffcient, r
P < 2.2e-16*
Time in million years

**Fig S8**

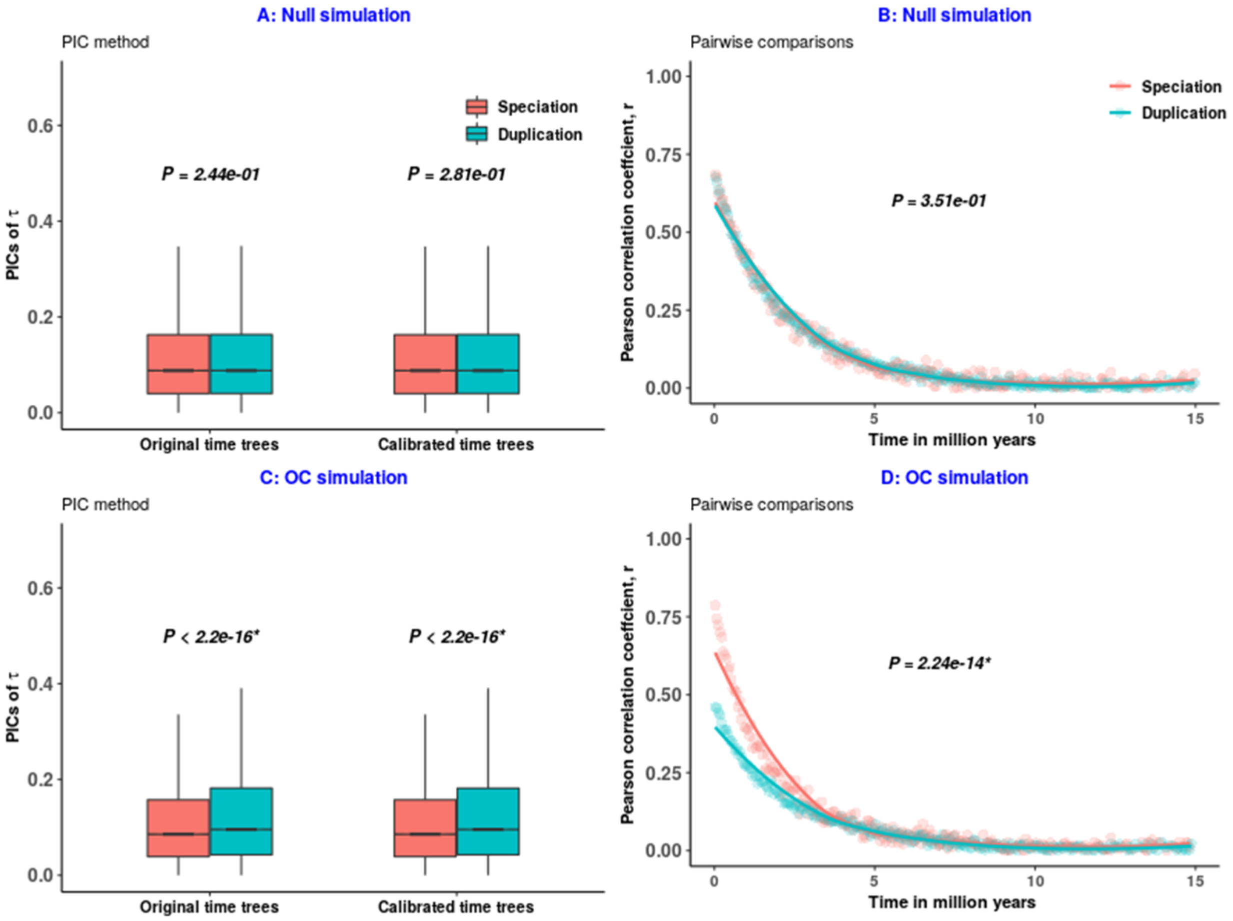
A: Null simulation
PIC method
Speciation
Duplication
P = 2.44e-01
P = 2.81e-01
PICs of τ
0.6
0.4
0.2
0.0
Original time trees
Calibrated time trees
B: Null simulation
Pairwise comparisons
Speciation
Duplication
P = 3.51e-01
Pearson correlation coeffcient, r
1.00
0.75
0.50
0.25
0.00
0
5
10
15
Time in million years
C: OC simulation
PIC method
P < 2.2e-16*
P < 2.2e-16*
PICs of τ
0.6
0.4
0.2
0.0
Original time trees
Calibrated time trees
D: OC simulation
Pairwise comparisons
P = 2.24e-14*
Pearson correlation coeffcient, r
1.00
0.75
0.50
0.25
0.00
0
5
10
15
Time in million years

**Fig S9**

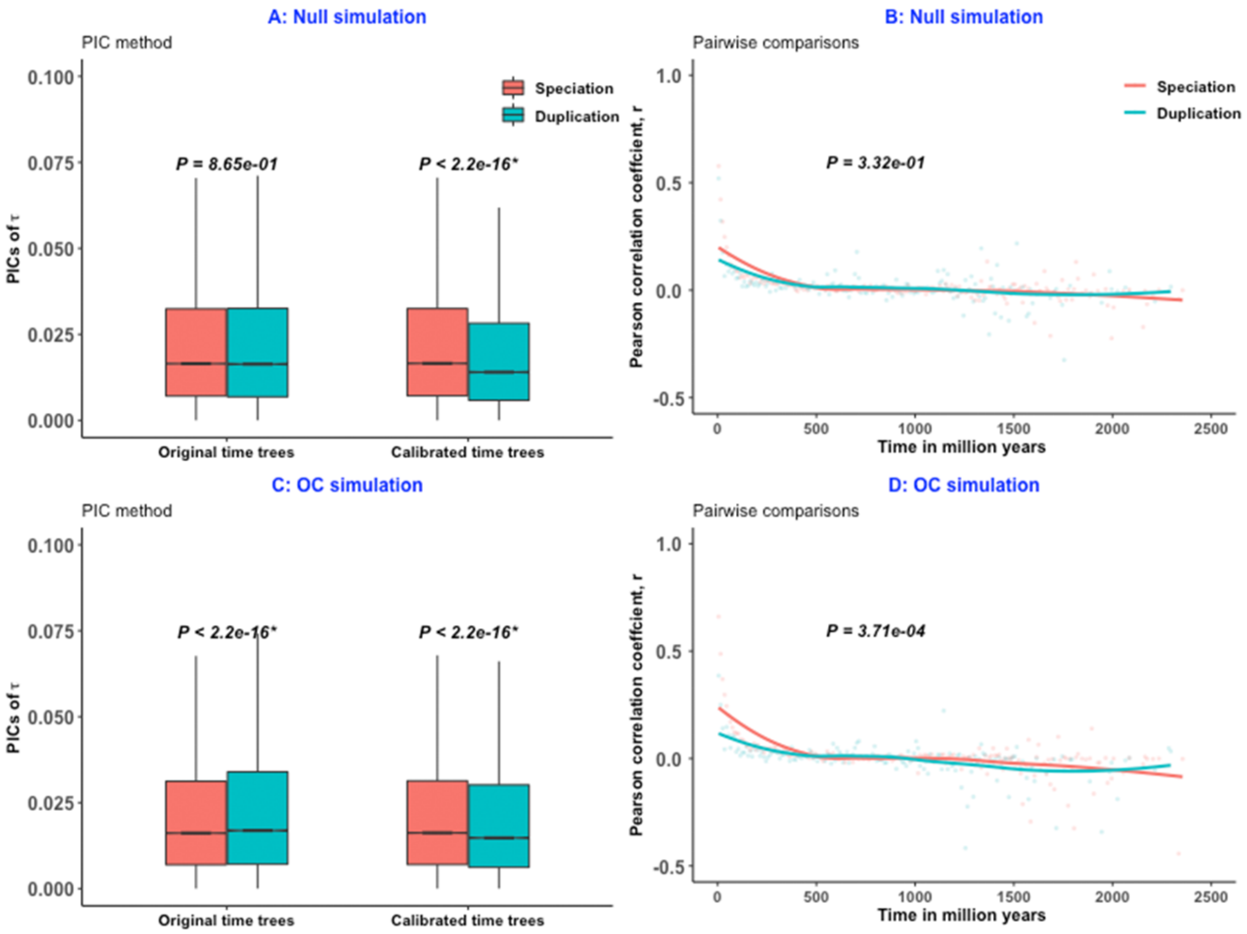
A: Null simulation
PIC method
PICs of τ
Speciation
Duplication
P = 8.65e-01
P < 2.2e-16*
Original time trees
Calibrated time trees
B: Null simulation
Pairwise comparisons
Pearson correlation coeffcient, r
P = 3.32e-01
Time in million years
C: OC simulation
PIC method
PICs of τ
P < 2.2e-16*
P < 2.2e-16*
Original time trees
Calibrated time trees
D: OC simulation
Pairwise comparisons
Pearson correlation coeffcient, r
P = 3.71e-04
Time in million years

**Fig S10**

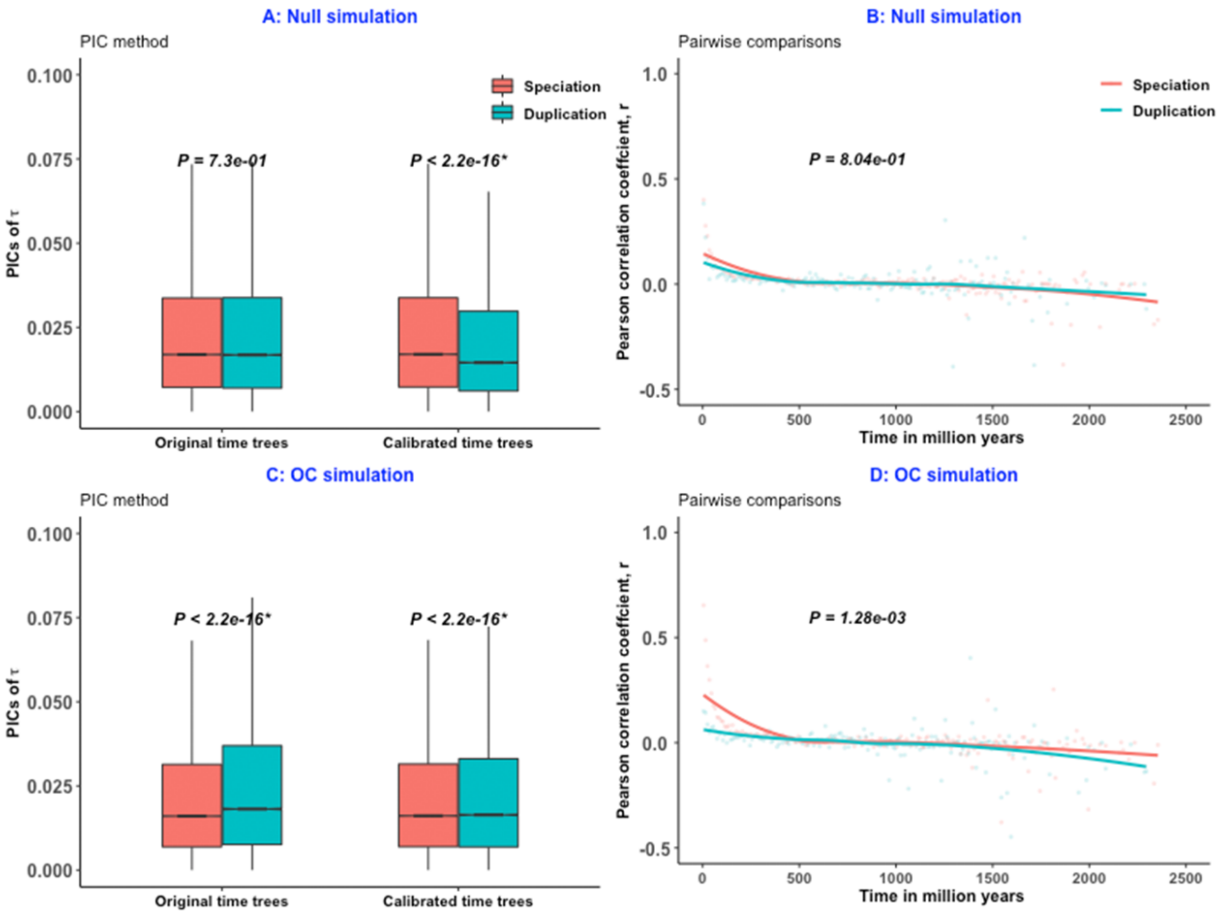
A: Null simulation
PIC method
Speciation
Duplication
P = 7.3e-01
P < 2.2e-16*
PICs of τ
Original time trees
Calibrated time trees
B: Null simulation
Pairwise comparisons
Speciation
Duplication
P = 8.04e-01
Pearson correlation coeffcient, r
Time in million years
C: OC simulation
PIC method
P < 2.2e-16*
P < 2.2e-16*
PICs of τ
Original time trees
Calibrated time trees
D: OC simulation
Pairwise comparisons
P = 1.28e-03
Pearson correlation coeffcient, r
Time in million years

**Fig S11**
